# Evaluating the ligands’ potency to modulate the fast inactivation of voltage-gated sodium channel

**DOI:** 10.1101/2025.07.21.665872

**Authors:** Beata Niklas, Milena Jankowska, Katarzyna Walczewska-Szewc, Bruno Lapied, Wiesław Nowak

## Abstract

Electrical impulse transmission along the nerve fiber in the form of the action potential is possible due to fast conformational changes of voltage-gated sodium channels (Na_v_) that control the sodium ions flow into the cell. The transition between functional states, called the gating mechanism, can be modulated by natural toxins and drugs. Here, we propose to use steered molecular dynamics (SMD) to investigate the ability of various ligands to impact the gating of *P. americana* cockroach Na_v_. By calculating mechanical forces required to relocate the inactivation particle to its binding pocket or to dislocate it, we assessed ligands’ efficacy in trapping a channel in a given state (open or fast inactivated). Importantly, we showed that sulfonamide PF-05089771 and phospholipid PIP2 act as insect Na_v_ channels inhibitors. We confirmed the ligands’ action by electrophysiological measurements of their ability to modulate the neural activity. Our approach, applied here on a cockroach channel, can be used in any other Na_v_, i.g, to evaluate new drug candidates.

## 1. Introduction

Electrical impulse transmission in excitatory cells is possible due to the rapid changes in membrane potential propagated along the nerve or muscle fiber as the action potential [1]. A key role in this process is played by voltage-gated sodium (Na_v_) channels, transmembrane proteins that undergo fast conformational changes to control sodium ion flow into the cell. In humans, nine subtypes of Na_v_ channels (hNa_v_1.1-hNa_v_1.9) are expressed, exhibiting distinct tissue distribution to fulfill unique physiological functions. In contrast, most insects carry only one sodium channel coding gene, relying on alternative splicing and RNA editing to generate variants with different pharmacological properties [2].

In the basic cycle, three main functional states are distinguished: resting (closed), open, and inactivated (see Fig. 1). The gating mechanism (transition between these states) is possible due to the channel structure [3]. The α- subunit, the core of a channel that conducts the flow of sodium ions, consists of a polypeptide chain of ∼2050 amino acids that folds into four homologous domains (DI–DIV) arranged in a domain-swapped configuration. Each domain is built by six transmembrane helices (S1–S6). Helices S5 and S6 form the ion-conducting pore with the two narrowest points in the selectivity filter and the activation gate. Four peripheral voltage-sensing domains (VSDs) comprise S1-S4 helices. When the change in membrane voltage reaches the threshold level, the positively charged S4 helices make the outward movement that leads to the opening of the channel and depolarization of the cell. It is followed by the fast inactivation mechanism to quickly (within 1-2 ms) block sodium ion flow. Upon repolarization, S4 helices move downward, the pore closes, and thus the channel comes back to the resting state.

**Figure 1.**
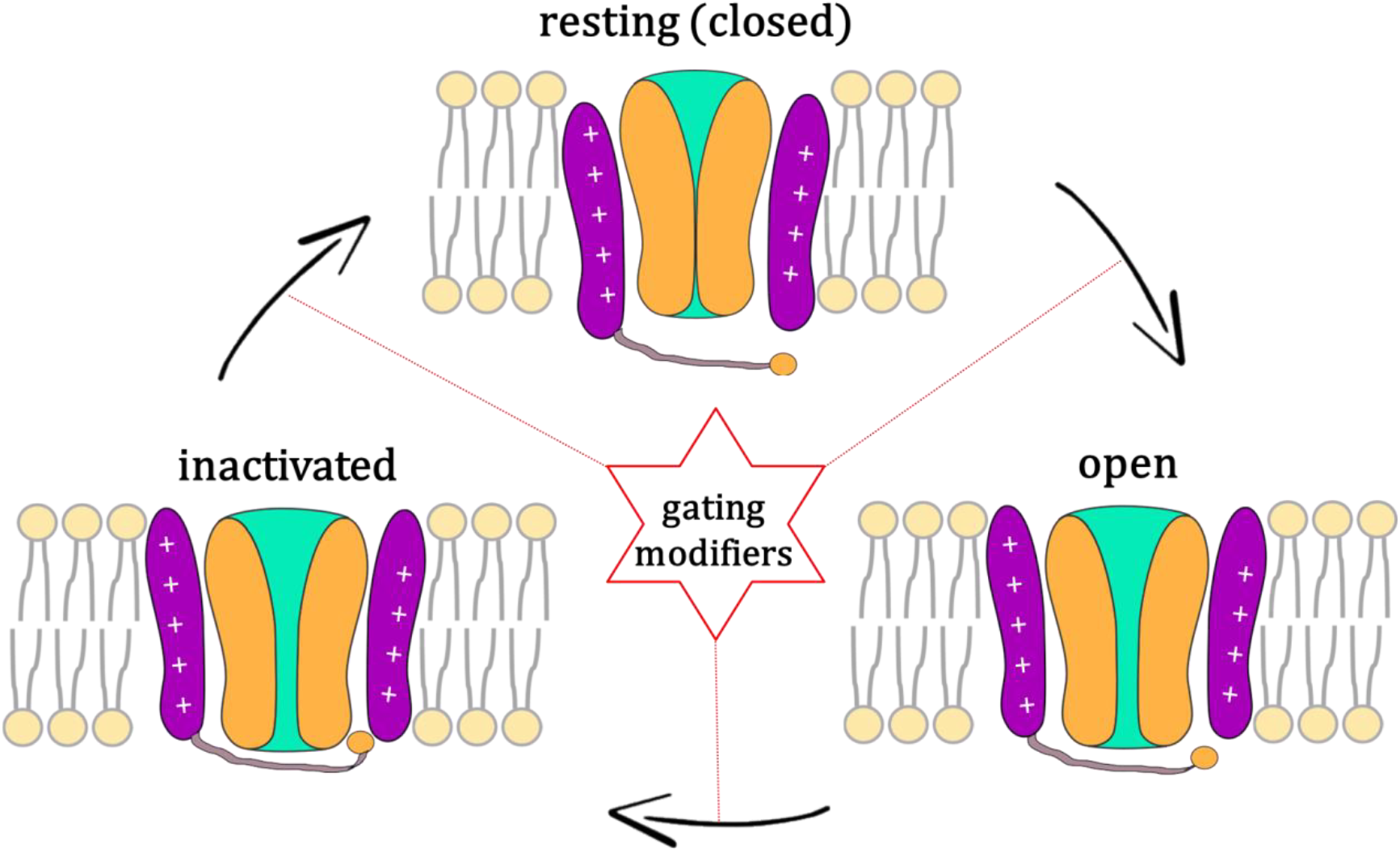
Schematic view on the gating cycle of voltage-gated sodium channel.

The gating mechanism, crucial for proper impulse transmission, is targeted by a wide range of naturally occurring toxins and small-molecule drugs that bind in distinct receptor sites, thus affecting the transition between states in various ways [4, 5]. Over the past decade, the search for new, selective therapeutics targeting mammalian Na_v_ channels has centered on channel areas that are less conserved among isoforms than the ion-conducting pore – VSDs. The greatest interest is in VSDIV since the movement of the S4-VSDIV helix triggers fast inactivation. It is naturally targeted by peptide toxins produced by scorpions and sea anemones [6, 7], which were extensively studied as drug or insecticide candidates. Although peptides exhibit high subtype selectivity and reduced off-target activity, good *in vitro* results do not translate into in vivo, possibly due to limited membrane permeability and lack of oral bioavailability [8]. In recent years, a considerable effort has been put into developing small-molecule compounds that would block S4- VSDIV movement, leading to the discovery of inhibitors with sulfonamide moieties. They exhibit high selectivity and have been investigated in clinical trials as potential pain modulators targeting hNav1.7 [8]. These compounds bound with high affinity to the extracellularly accessible site on VSDIV, effectively trapping it in the activated (“up”) state, allowing the channel to undergo inactivation but preventing a return to a resting state, which blocks the sodium ion conductance [9]. Despite excellent in vitro results, these compounds have not passed any clinical trials to date, most likely due to insufficient concentration at the binding site. They were, however, well tolerated, with no serious adverse event observed, which encourages searching for alternative uses.

Although one of the most efficient ways to modify channel gating is to impair the fast inactivation, this process is still not fully understood. Numerous studies in the past described the inactivation gate as a part of the cytoplasmic linker connecting DIII and DIV (DIII-DIV linker, dark blue in Fig. 2), with the crucial region centered on the Ile-Phe-Met (IFM) motif in mammals and Met-Phe-Met (MFM) in insects [10]. For a long time, the hinge-lid model, in which the hydrophobic IFM fragment, called inactivation particle, physically occludes the intracellular mouth of the pore to stop sodium ion flow, was accepted [10-12]. However, the emergence of many high-resolution cryo-electron microscopy (cryo-EM) structures of inactivated-state mammalian Na_v_s, where IFM is not located in the pore axis, as well as more recent structural studies, has led to a reinterpretation of the fast inactivation process [13-16]. Currently, it is known that binding of IFM to its pocket triggers an allosteric pathway of conformational changes involving displacement of the DIV S6 helix towards the ion permeation pathway that results in partial block of sodium conductance [17]. Thus, the IFM motif, historically called the inactivation gate, is not a gate itself. The allosteric pathway between IFM binding and residues of S6 helices that occlude the pore, as well as molecular detail of the final step, are still elusive [17] and currently extensively studied [18].

**Figure 2.**
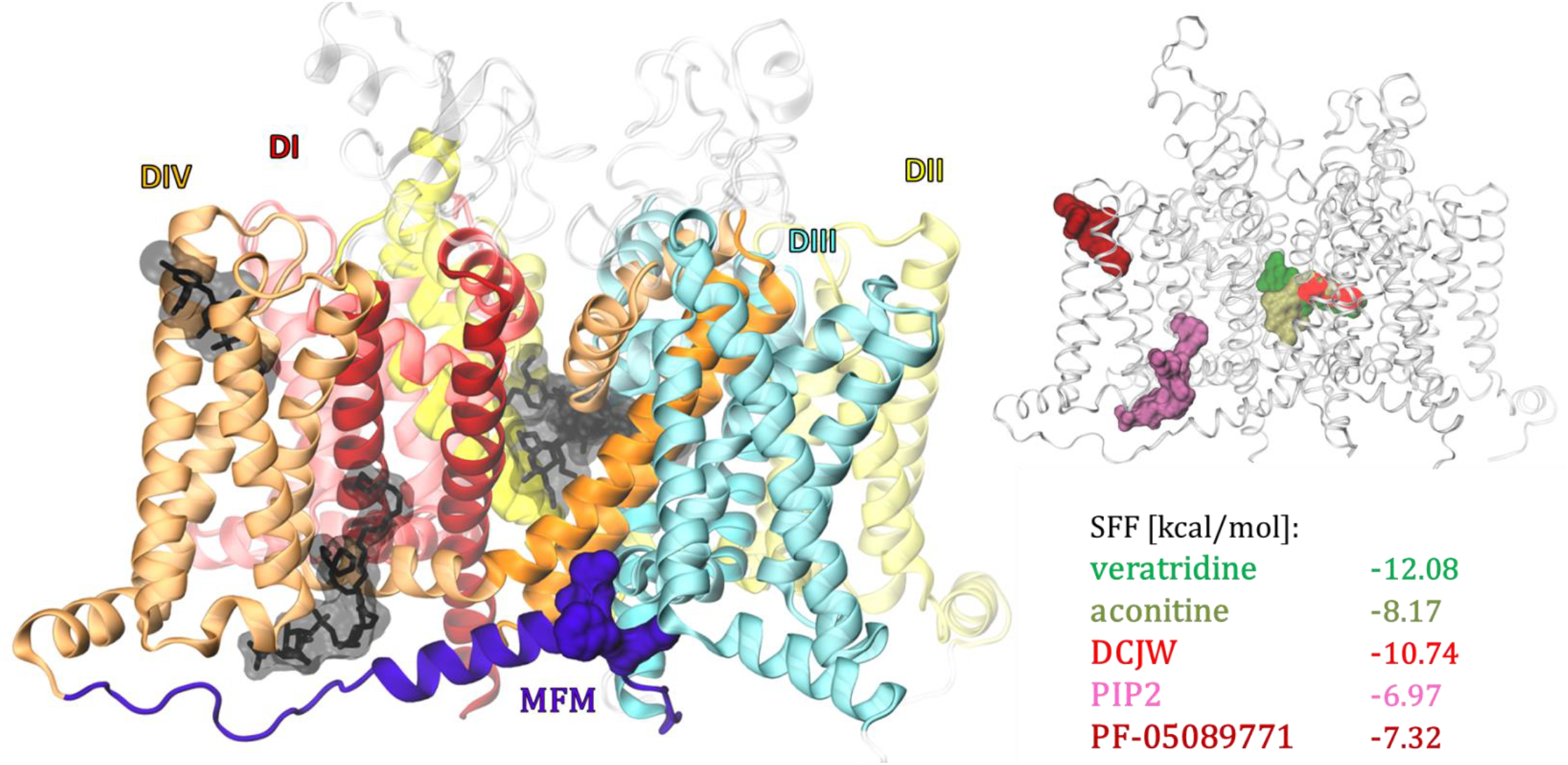
Investigated ligands docking to the insect voltage-gated sodium channel. Left panel shows the structure of the open state PaNav1 model, with four domains colored: DI – red, DII – yellow, DIII – blue, DIV – orange. The DIII-DIV linker is in violet with the inactivation particle MFM motif shown in a surface representation. Ligands are shown in black. In the right panel, the lowest energy position of ligands, together with their affinity to the channel, measured by a smina scoring function (SSF), are in colors (veratridine – dark green, aconitine – olive, DCJW – red, PIP2 – pink, and PF-05089771 – dark red). The detailed views of interactions of all ligands are shown in SI Fig. 4-8.

Molecular Dynamics (MD) is an extensively used *in silico* technique that predicts the time evolution of interacting atoms, serving as a computational microscope in molecular biology [19]. It plays an important and continuously increasing role in the quest to connect the structure and function of biomolecules. However, the use of classical MD is limited by the timescale constraints, i.e., the inability to efficiently sample rare events and slow processes. Steered molecular dynamics (SMD) addresses this limitation by applying time-dependent external forces to induce specific transitions, e.g., conformational changes in protein. Moreover, the calculation of forces required for a transition to happen allows for comparison of the impact of external factors like the presence of ligands.

Here, we applied SMD, together with extracellular recording of action potentials, to investigate the ability of various ligands to modify gating of cockroach Na_v_. By calculating mechanical forces required to relocate the MFM particle to its binding pocket (from the open structure) or to dislocate it (from the inactivated-state channel) in an unliganded (apo) versus ligand-bound channel, we are able to compare ligands’ efficacy to trap the channel in a given state. We validated the *in silico* results with electrophysiological measurements of neural activity on a model organism, Periplaneta americana. Our simple approach, applied here on a cockroach channel, can be used in any other Na_v_, i.g, to evaluate new drug candidates.

## 2. Materials and methods

### 2.1 Systems preparation

The homology models of an α subunit of the Periplaneta americana sodium channel protein PaNav1 in its open, inactivated, and closed states were built using the SWISS-MODEL server [20], using rat rNav1.5 (PDB code: 7FBS[21]), human hNav1.2 (PDB code: 6J8E[22]), and cockroach NavPas (PDB code: 8VQC[23]) experimental structures, respectively, as templates. Models were built based on the D0E0C1_PERAM amino acid sequence provided by the UniProtKB database [24]. Two disordered intracellular loops (between DI-DII and DII-DIII), absent in any sodium channel protein experimental structure, were removed from each model. The protein preparation module of Schrodinger Maestro [25] was used to assign protonation states, add hydrogen atoms, and minimize all three models, as well as to introduce the N1739D substitution and further minimize the mutated open-state model. Molecular docking of all ligands was performed using smina package [26], a fork of Autodock Vina [27]. Ten independent runs (giving up to 100 poses each) of flexible ligands docking to each model were conducted with default parameters, followed by a minimization of the best-scored poses. Inputs for equilibrium MD simulations were generated using CHARMM-GUI membrane builder [28-30]. A heterogeneous bilayer model composed of 400 lipid molecules in proportions: 38% DOPE, 18% DOPS, 16% DOPC, 13% POPI, 11% SM (CER180), 3% DOPG, and 1% PALO 16:1 fatty acid was built to mimic an insect-like membrane, as proposed for a fruit fly Drosophila melanogaster mAChR-A GPCR [31], and a Na_v_ of Anopheles gambiae mosquito [32]. Water molecules in the TIP3P model were added above and below the lipids to generate a 20 Å thickness layer further neutralized with Na^+^ and Cl^-^ ions to the final concentration of 0.15 M. Six steps of equilibration composed of NVT dynamics followed by NPT dynamics with gradually decreased restraint force constants to various components were run using NAMD 2.14 [33] with the CHARMM36 force field applied. A 1 ns cMD simulation was performed for each system before SMD.

### 2.2 SMD simulations

Inputs for SMD were generated based on cMD outputs, using multiSMD – a Python toolkit designed to measure the disruption forces of proteins or protein domains by measuring forces required to pull them away from each other. The original script (that can be found at https://github.com/kszewc/multiSMD) calculates the centers of masses (COMs) of two selections and creates a set of vectors corresponding to the pulling directions. We modified a multiSMD method for multidirectional SMD to suit the goals of our project. Modifications included:

- to mimic fast inactivation and calculate the forces required to move the MFM particle to its binding pocket, we redirected the generated set of vectors. Instead of extracting domains, we simulated the insertion of the MFM particle into its binding pocket.
- We used two separate selections: one for fixing the atoms (‘protein and name CA and resid 387 972 1498 1791’), and the second one for generating the principal axis of pulling, which serves as a vector connecting the centers of mass of the inactivation particle MFM motif (‘protein and name CA and resid 1565:1567’) and its binding pocket (here: ‘protein and name CA and resid 1411 1552 1555 1739 1740 1842 1850’). Fixing the atoms of the selectivity filter enabled us to distinguish fast inactivation from the slow one, as the selectivity filter is engaged only in the latter (RMSD between fast-inactivated and open states is equal to 0.3 Å).
- We modified theta and phi angles to focus the sampling on the path linking MFM with its pocket. We thus investigated nine directions of pulling: theta (0^°^, 150^°^, 165^°^) and phi (0^°^, 90^°^, 180^°^, 270^°^)

After 1 ns cMD, we run 20 ns of SMD with a timestep of 1 fs in a NVT ensemble so that the pulling force is adjusted to provide a constant velocity in the direction of pulling. To select the best SMD direction, we visually analyzed the trajectories (checking if MFM moves to its pocket) and compared the forces required in simulations of the unbounded channel (apo) with ligand-bound. We started with ligands with known effects on Na_v_s: DCJW (which promotes inactivation) and veratridine (which inhibits inactivation) and found the best pulling direction to be theta=150^°^, phi=180^°^. We used this direction to investigate the effects of other ligands (3×20 ns for each ligand).

In simulations of the recovery from fast inactivation, we changed again the orientation of vectors and tested the following directions of pulling: theta (0^°^, 30^°^, 60^°^) and phi (0^°^, 90^°^, 180^°^, 270^°^). The most appropriate direction, theta=60^°^, phi=90^°^, was used further in simulations with ligands (3×20 ns for each system).

Hydrogen bonds were calculated using the MDAnalysis Python package [34] with the binding pocket residues and F1566 as a selection.

### 2.3 Electrophysiology

Experiments were conducted on male American cockroaches, *Periplaneta americana*. The insects were bred and maintained under appropriate conditions at Nicolaus Copernicus University at a temperature of 27–29°C, with unlimited access to water and food (cat food, oatmeal, and apples). To allow adaptation, cockroaches were transferred from the breeding facility to the procedure room 24 hours before the experiment.

According to European Union regulations, studies on invertebrates do not require approval from an ethics committee; however, all procedures were carried out by the 3R principle.

Electrophysiological recordings were performed on cercal nerves containing sensory neurons originating from the cerci appendages. For this purpose, the dorsal part of the abdomen was dissected, and the digestive system, along with accessory glands, were removed to expose the terminal abdominal ganglion, the cercal nerves, and a section of the connective nerve leaving the ganglion. The preparation was mounted on a plate and continuously irrigated with physiological saline (pH = 7.4; 210 mM NaCl, 3.1 mM KCl, 5 mM CaCl_2_, 5.4 mM MgCl_2_, 5 mM Hepes). An extracellular recording electrode (with a typical DC resistance of 0.3 V and an impedance of 20–30 kΩ at 1 kHz) was placed on the cercal nerve, while a reference electrode was positioned in the physiological saline at the edge of the preparation. Bioelectric signals were recorded using an open-source amplifier [35] and visualized on a computer with open-source software Spike Recorder (https://github.com/BackyardBrains/Spike-Recorder).

During recordings, the left cercus was stimulated by air puffs at a frequency of 0.75 Hz, generated by a mechanostimulator. The response to stimulation, observed as a brief and rapid increase in nerve activity, was used to confirm the preparation’s condition. For analysis, the frequency of spontaneously generated action potentials was evaluated.

Veratridine (Merck, Poland), aconitine (Merck, Poland), indoxacarb (Merck, Poland), and PF-05089771 (Merck, Poland) were initially dissolved in DMSO at a concentration of 10 mM and then diluted in physiological saline to a final concentration of 10 µM. PIP2 in a chloroform/methanol solution (phosphatidylinositol 4,5-bisphosphate, Merck, Poland) was air-dried, and the remaining film was dissolved in DMSO to obtain a concentration of 10 mM before being diluted in physiological saline to a final concentration of 10 µM. All tested solutions were applied near the cercal nerve in a 20 µL volume when the frequency of spontaneous action potentials remained stable for at least 2 minutes but not earlier than 5 minutes before the start of recording.

## 3. Results and discussion

### 3.1 Homology modeling

We built three homology-based models of the cockroach PaNa_v_1 channel using experimental structures of eukaryotic Na_v_s representing open, inactivated, and resting states as templates (PDB codes: 7FBS[21], 6J8E[22], and 8VQC[23], respectively).

We ensured that in the open-state model, the S4 VSDIV is in activated (“up”) conformation, which is necessary for the inactivation to begin according to the sliding helix model. In this model, the S4 gating charges move outward through the hydrophobic constriction site (HCS) and exchange ion pair interactions with the intracellular negative cluster (INC) and extracellular negative cluster (ENC) of negatively charged amino acid residues [36]. In both our open- and inactivated-state models, four gating charges (R1703, R1706, K1709, R1712) are on the extracellular side of the HCS, while the fifth and sixth (R1715 and R1718) are on the intracellular side, interacting with the INC (SI Fig. 1). In contrast, in the resting-state model, only two gating charges (R1703, R1706) are above HCS.

The main structural differences between the open- and inactivated-state models are centered on the DIII-DIV linker containing the inactivation particle MFM motif, the intracellular ends of S6 helices (the activation gate region), and the intracellular beginnings of the VSD domains (red-colored in SI Fig. 2).

### 3.2 Ligands and docking

Ligands targeting Na_v_ channels can be divided into blockers and gating modifiers. While blockers (e.g., tetrodotoxin) physically reduce the sodium ion flow through the channel, gating modifiers affect action potential by disrupting the transition between main conformational states (see Fig. 1). The latter can be further divided into activators and inhibitors. Activators increase sodium ion flow by keeping the channel open for longer, which is achieved by slowing down/disrupting the fast inactivation process. Inhibitors may act by promoting fast inactivation and/or inhibiting the recovery from fast inactivation (i.e., returning to the resting state so that the channel is available for another depolarization).

We selected ligands with known activity on the insect Na_v_ to validate the method and then proceed to investigate the mode of action of ligands with not fully understood effect on channel gating (Table 1).

**Table 1.**
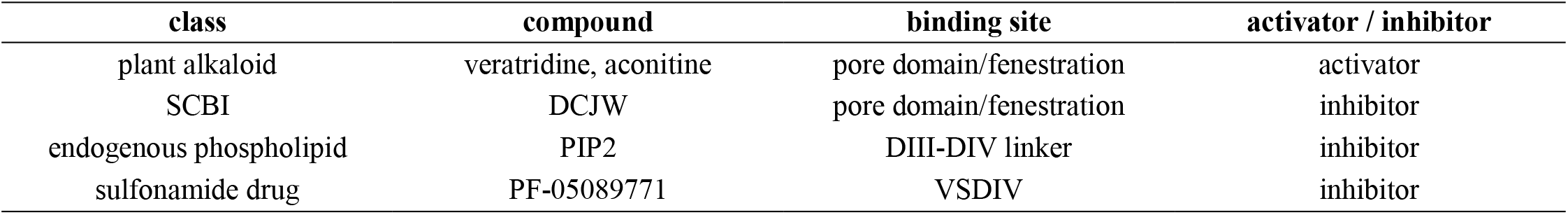
Ligands investigated.

| class | compound | binding site | activator / inhibitor |
| --- | --- | --- | --- |
| plant alkaloid | veratridine, aconitine | pore domain/fenestration | activator |
| SCBI | DCJW | pore domain/fenestration | inhibitor |
| endogenous phospholipid | PIP2 | DIII-DIV linker | inhibitor |
| sulfonamide drug | PF-05089771 | VSDIV | inhibitor |

**Table 1.** Minimal distance between centers of masses of MFM particle and its binding pocket observed during SMD trajectories.

| system | apo | N1739D | aconitine | veratridine | DCJW | PF-05089771 | PIP2 |
| --- | --- | --- | --- | --- | --- | --- | --- |
| minimal distance [Å] | 5.03 | 10.18 | 8.35 | 8.24 | 4.60 | 4.11 | 6.09 |

Veratrum and Aconitum plants were considered a potential source of insecticidal compounds over the last century, as they contain highly toxic steroidal alkaloids (e.g., veratridine, aconitine) that activate Na_v_ channels. However, these compounds are non-selective towards arthropods and can cause rapid cardiac failure in humans. Veratridine is currently used in science as a potent gating modulator that opens the channel and traps it in the conducting state [37, 38]. Aconitine-modified channels inactivate completely, although with slower kinetics [39]. We used these toxins as known channel activators.

DCJW is a neurotoxic derivative of a prodrug indoxacarb, a member of the sodium channel blocker insecticide (SCBI) class, which are potent Na_v_ channel inhibitors. Indoxacarb is metabolized to more potent DCJW through decarbomethoxylation in arthropods but not in mammals, which makes this compound selective towards insects and has led to its widespread use in agriculture. DCJW reaches its binding site in the ion-conducting pore through one of the hydrophobic tunnels that link the plasma membrane with the central cavity of the channel, called fenestrations [32]. Although SCBIs are called blockers, they modify gating by trapping Na_v_ channels in a non-conducting state, and thus can be classified as inhibitors.

Phosphatidylinositol 4,5-bisphosphate (PIP2), a phospholipid component of the inner leaflet of cell membranes, is an important signaling molecule not only as a substrate for second messenger generation but also as a direct modulator of membrane proteins [40]. PIP2 is known to interact with G protein-coupled receptors and numerous ion channels, including voltage-gated, inwardly rectifying and calcium-activated potassium channel families, voltage-gated calcium channels, calcium-activated chloride channels, transient receptor potential channels, cyclic nucleotide-gated channels, purinergic-regulated P2X channels and epithelial sodium channels [41]. This phospholipid acts as the allosteric modulator of channel activity with positive or negative effects on ion conductance, depending on the channel type. Recently, PIP2 was also shown to reduce Na_v_1.4 current by regulating the voltage-dependence of activation, accelerating the transition to the inactivated state, and slowing recovery from inactivation [42]. Further MD studies revealed PIP2 binding to the conserved site within the DIII–DIV linker of inactivated-state Na_v_1.4 and Na_v_1.7, which slows the coupling between VSD and the pore [43]. We proceed to investigate if PIP2 can also modulate insect Na_v_, embedded in an insect-like membrane model.

Finally, encouraged by the lack of serious adverse events found in clinical trials for the treatment of acute or chronic pain, we tested if PF-05089771, known as a selective sulfonamide inhibitor targeting hNa_v_1.7, can also modify insect neuronal conductance.

We started by docking all described ligands (see SI Figure 3 for their chemical structures) to the open state PaNa_v_1 model. In their lowest energy poses, veratridine, aconitine, and DCJW occupy the pore domain and fenestrations, which agrees with previous studies [32, 44-46]. PIP2 binds in the cavity formed by the DIII-DIV linker, VSD helices S3-S4 from DIV, and S4-S5 linker from DIV, as recently described [43]. PF-05089771 binds in the extracellular part of VSD from DIV to interact with the gating charges of the S4 helix, like in a cryo-EM structure of hNav1.7 [46]. Ligands in their lowest energy poses are shown in Figure 2, and the channel-ligand interactions are presented in SI figures 4-8.

We also tested how the mutation of N1739, which was found to inhibit fast inactivation in the Nav1.2 channel (N1662A in rat Nav1.2 [47] and N1662D in human Nav1.2 [48]), affects forces required for MFM binding. N1739, located in the cytoplasmic end of the DIV S5 helix, forms a hydrogen bond with Q1571 of the DIII-DIV linker, thus stabilizing the MFM motif in its bound state. The N1662D substitution in the *SCN2A* gene coding hNav1.2 is associated with early-infantile developmental and epileptic encephalopathy.

### 3.3 Multidirectional SMD to mimic the motion of the MFM particle

During fast inactivation, the IFM particle of DIII-DIV linker forms a hydrophobic interaction with its receptor site, for a long time expected to be located in the intracellular part of the ion-conducting pore. The actual location of the IFM binding pocket was revealed when the cryo-EM structures representing inactivated-state Na_v_ channels emerged. IFM binds to the hydrophobic pocket formed by the S4-S5 linkers of DIII and DIV, and the S6-DIV helix, adjacent to the intracellular activation gate (see SI Fig. 9). Notably, this receptor site only forms when the VSDIV is in activated “up” conformation (see SI Fig. 1).

Knowing the amino acid residues forming mammalian IFM or insect MFM binding pocket enabled us to define the principal axis of pulling in SMD to “push” the inactivation particle to its pocket, starting from the open state channel. This principal axis of pulling is defined as a vector connecting the centers of mass of the MFM particle and residues creating its binding pocket. Since selection of pulling direction is one of the crucial parameters in SMD simulations [49], we started by performing SMD in multiple directions, i.e. using a set of vectors, characterized by small variations in theta (0^°^, 150^°^, 165^°^) and phi (0^°^, 90^°^, 180^°^, 270^°^) angles within spherical coordinates. Each sampling vector represents a unique direction for pulling, giving nine independent SMD simulations (SI Fig. 10a).

From SMD outputs, we calculated the dependence of pulling force on the distance between the centers of masses of MFM and its binding pocket, which allowed us to observe the system’s response to the applied force directly. Based on distance vs. force profiles of unliganded Na_v_ channel (apo) and with DCJW- or veratridine-bound, as well as the visual inspection (trajectory analysis), we have chosen θ=150^°^, φ=180^°^ as the best pulling direction for SMD modeling of fast inactivation (see SI Fig. 11 for all directions, and Fig. 3 for the best one).

**Figure 3.**
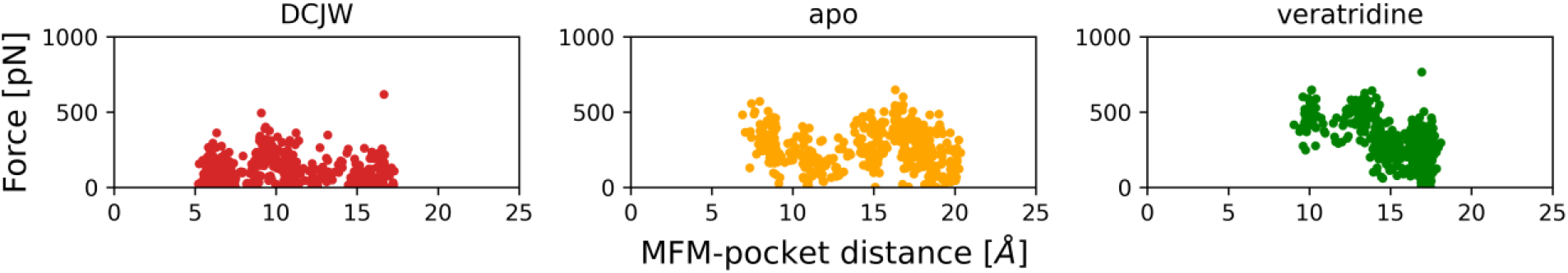
The force vs. distance plots from SMD simulation in the θ=150^°^, φ=180^°^ direction from the main pulling axis for apo (orange), DCJW-bound (red), and veratridine-bound Na_v_ reflecting the ligands’ ability to affect the MFM particle movement to its binding pocket.

As shown in Fig. 3, the MFM particle of the DCJW-bound channel moved the closest to its binding pocket (with the minimal value of 5.2 Å, when compared to 6.9 Å for apo, and 9 Å for veratridine-bound Na_v_ channel). Even though simulation with veratridine required the highest force (with the mean value of 309 pN, when compared to 261 pN for apo, and 128 pN for DCJW), the MFM particle could not reach its pocket. This reflects, as expected, that the fast inactivation is facilitated with DCJW bound and hampered in the presence of veratridine.

As inhibitors may act by enhancing fast inactivation and/or trapping the channel in the inactivated state, we proceed to analyze their impact on the recovery from inactivation. With no prior knowledge of ligands effect, we tested the method on the ligand-free channel. Here, we started SMD from the inactivated-state model to pull the MFM particle out of its pocket. We tested nine pulling directions: theta (0^°^, 30^°^, 60^°^) and phi (0^°^, 90^°^, 180^°^, 270^°^) (SI Fig. 10b). Apart of the distance vs. force profiles (SI Fig. 12, middle panel), we also compared the change in pulling force over time (SI Fig. 12, left panel), and the number of hydrogen bonds formed between F1566 (from MFM motif) and the MFM binding pocket (SI Fig. 12, right panel). For simulations with inhibitors, we have chosen θ=60^°^, φ=90^°^as the best pulling direction for SMD of recovery from fast inactivation, which we discuss further.

Fig. 4 shows the schematic view of MFM displacement in SMD simulations of fast inactivation (left panel), and recovery from fast inactivation (right panel) in chosen pulling directions.

**Figure 4.**
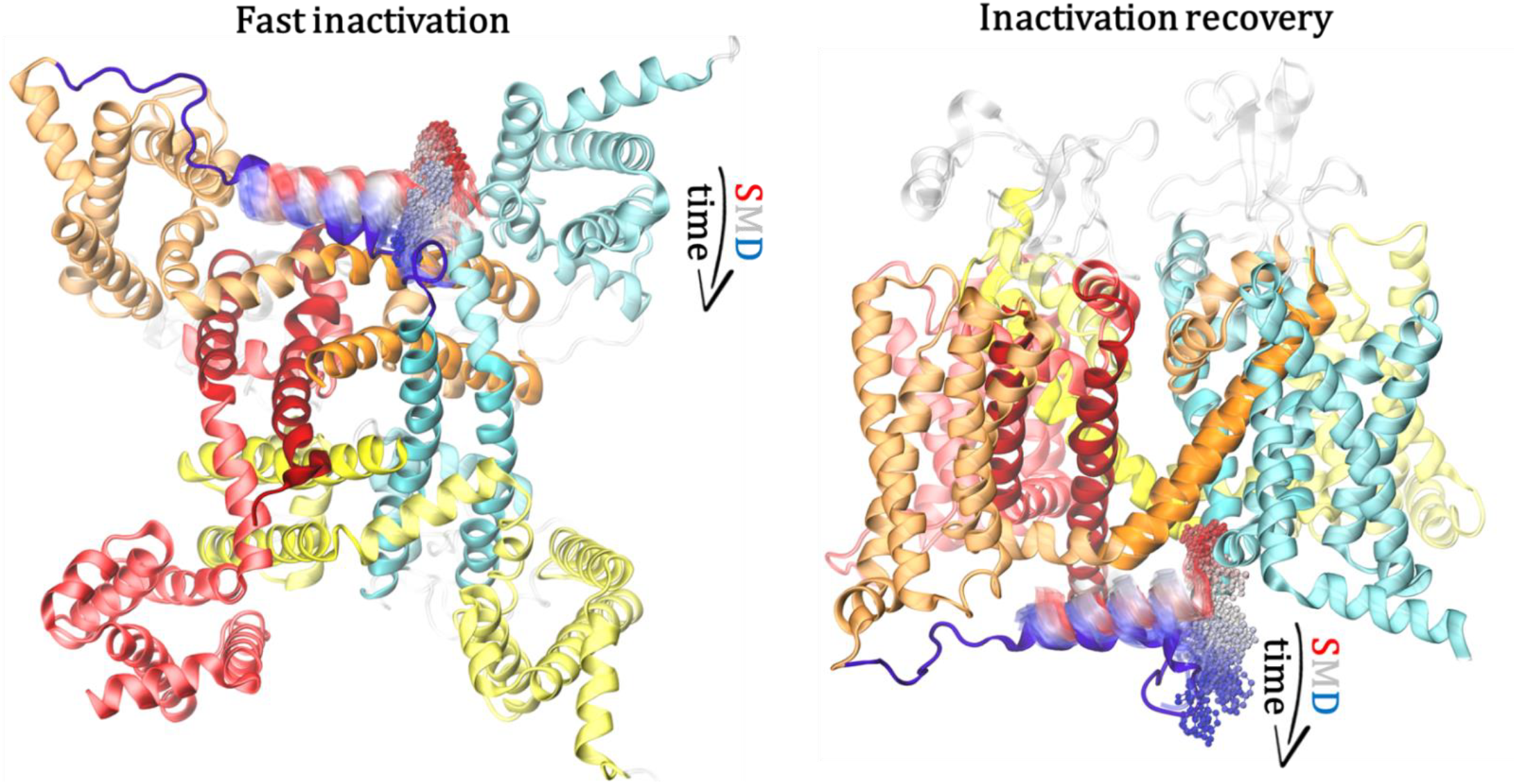
Scheme of SMD simulation representing fast inactivation (left, bottom view) and the recovery from fast inactivation (right, side view) for the best pulling directions (θ=150^°^, φ=180^°^, and θ=60^°^, φ=90^°^, respectively). The displacement of F1566 (dots) and the whole DIII-DIV linker (cartoon) during the 20 ns-long SMD simulation of unliganded channel is shown, where the beginning of the trajectory is in red, middle in white, and the end in blue.

### 3.4 SMD of MFM binding – mimicking the first stage of fast inactivation

Having the best direction of pulling chosen, we proceed to investigate the effect of other ligands, for some of which the impact on insect channel gating is unknown (three SMD runs for each). As the SMD simulation as short as 20 ns proved to be sufficient to observe the MFM binding in the unliganded channel, we kept this value for all further SMD runs. The force-time and force-distance profiles, together with hydrogen bonds analyses for all investigated ligands are presented in Fig. 5.

**Figure 5.**
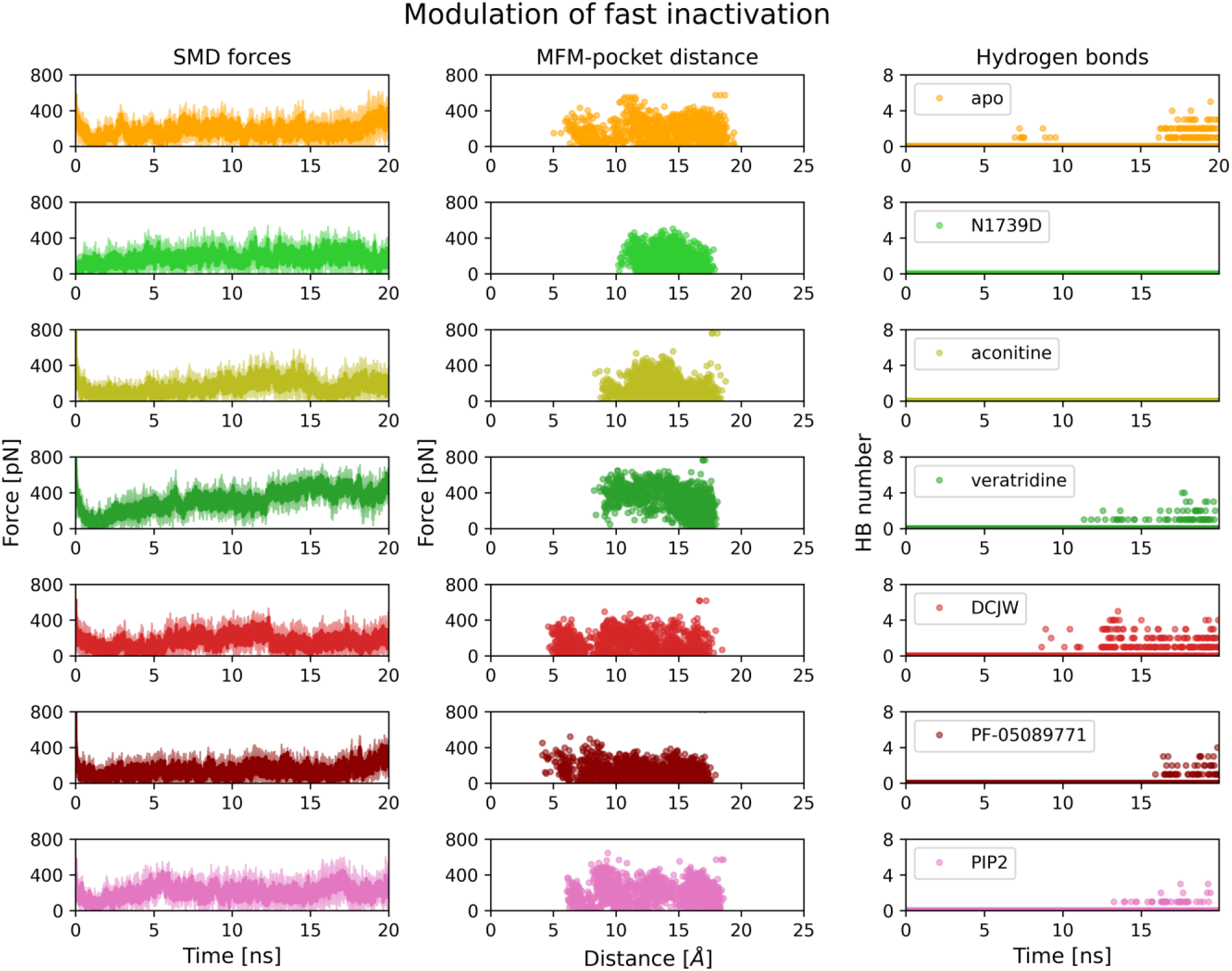
The ligands impact on the fast inactivation process of insect voltage-gated sodium channel. The left panel represents the change in forces required to push the MFM particle to its binding pocket over time. The middle panel shows the change in force corresponding to the distance between centers of masses of the MFM particle and its’ binding pocket. On the right panel, the number of hydrogen bonds between F1566 and the pocket is shown. Unliganded (apo) channel is shown in orange, channel with the ligands and the mutation that inhibit fast inactivation are in green (N1739D mutant in light green, aconitine in olive, and veratridine in dark green), followed by channel with presumed blockers (DCJW in red, PF-05089771 in dark red, and PIP2 in pink). Data are based on 3×20 ns-long SMD simulations for each case.

The highest forces were required in simulations with a potent Na_v_ activator, veratridine (Fig. 5, dark green), which indicates that inactivation was the most impeded in the presence of this ligand. Although hydrogen bond formation was observed, these were due to the interactions at the entrance to the binding pocket, where F1566 stuck until the end of one trajectory. The minimal distance between COMs of the MFM particle and its binding pocket was very similar for veratridine and aconitine, however, the difference in mean forces (311.4±4.6 pN and 153.3±16.7 pN, respectively) indicates higher potency of veratridine. Fast inactivation was also blocked in the mutated channel, although one should note that the N1739D substitution itself decreases the tracked distance as N1739/D1739 contributes to the MFM binding pocket.

In contrast, the full binding of the MFM particle to its pocket was observed in simulations of unliganded (Fig.5, apo, orange) and inhibitor-bound channels (red, dark red, and pink) with comparable force profiles. However, their differences were too subtle to conclude that tested inhibitors promote fast inactivation. Thus, the method proposed here might be more effective in estimating the protein’s resistance to conformational change (and hence the ligand’s effectiveness in holding the channel in a given conformation) rather than favoring the transition between the functional states of Na_v_.

### 3.5 The recovery from fast inactivation SMD

As the effect of inhibitors in promoting fast inactivation was subtle, we proceeded to investigate how their presence affects the recovery from fast inactivation (Fig. 6). We thus docked ligands to the inactivated-state PaNa_v_1 model and ran SMD of ligand-bound Na_v_, pushing MFM motif from its binding pocket. The activators were not tested here due to the state-dependence of their binding to the open channel [50].

**Figure 6.**
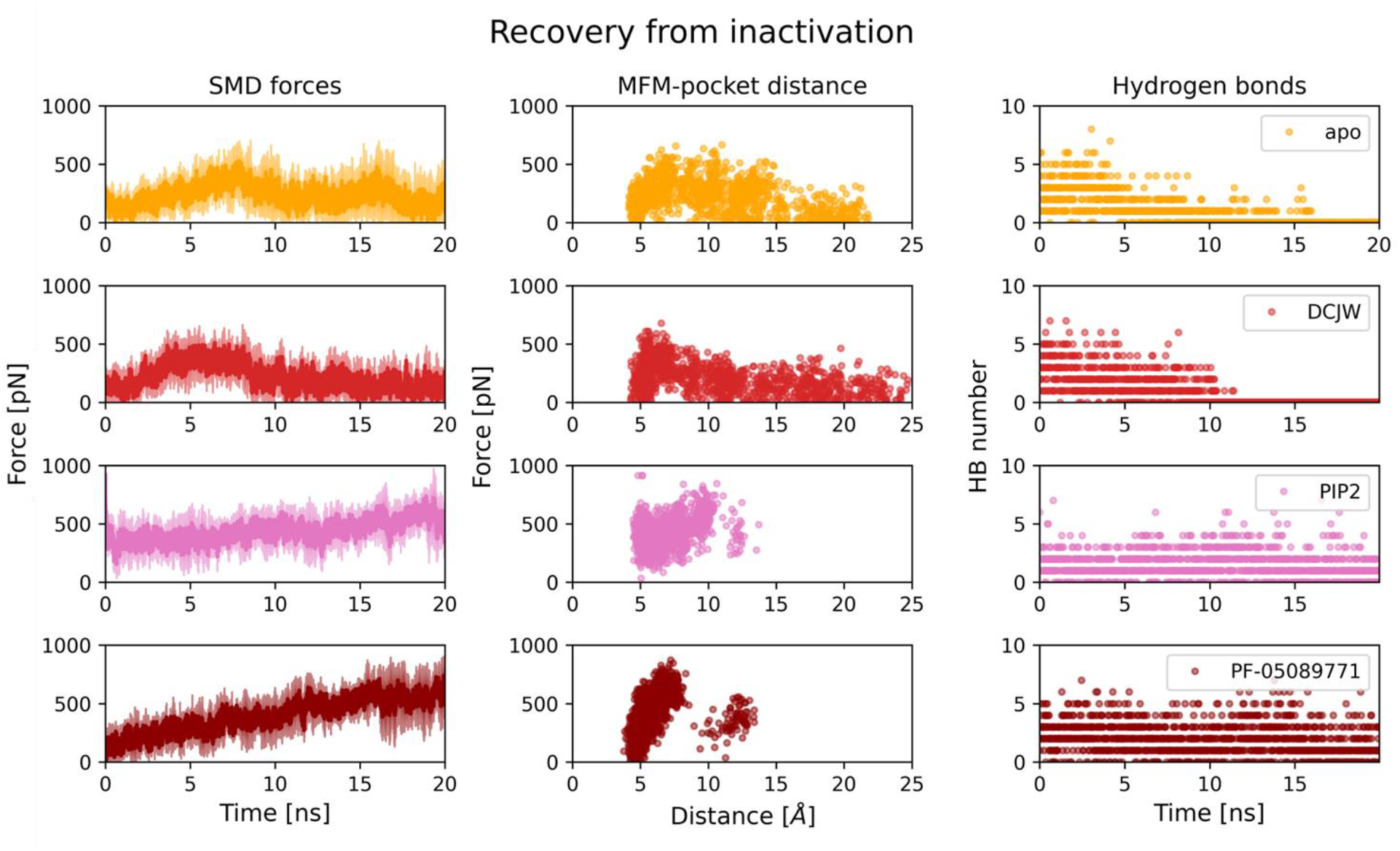
The ligands impact on the recovery from fast inactivation of insect voltage-gated sodium channel. Left panel represents the change in forces required to pull the MFM particle from its binding pocket over time. The middle panel shows the change in force corresponding to the distance between the centers of masses of the MFM particle and its’ binding pocket. On the right panel, the disruption of hydrogen bonds between F1566 and the pocket is shown. Unliganded (apo) channel is shown in orange, DCJW- bound in red, PIP2-bound in pink, and PF-05089771-bound in dark red. Data are based on 3×20 ns-long SMD simulations for each ligand.

Surprisingly, the forces required to pull MFM from its binding pocket in the DCJW-bound Na_v_ channel are nearly the same as for the apo channel. This may indicate, as previously reported [51, 52], that DCJW interacts with the channel in a slow-inactivated state, which cannot be investigated by the method used. There is, however, a striking difference between the unliganded channel and with other inhibitors bound. In simulations with PIP2 and PF- 05089771, the forces required to pull out the MFM motif are constantly increasing, and even the value of 800 pN is not enough to displace is by 10 Å. Also, the number of hydrogen bonds between F1566 and the MFM pocket is constant when PIP2 or PF-05089771 is bound. It is thus clear that these two ligands (acting *via* very different binding sites) prolong the recovery from fast inactivation.

In our analysis, we found that Na_v_ gating can also be tracked by measuring the frequency of F1565-F1552 interactions. As F1552, a part of DIII S6, lies deepest in the MFM binding pocket, the F1565-F1552 is the longest distance from the open-state starting point. Thus, this contact is the last to occur in fast inactivation and the first to be lost in the recovery from fast inactivation. Interestingly, the F1565-F1552 contact did not occur in any SMD replica with any of the activators, but it emerged at the end of apo and inhibitors-bound Na_v_ simulations (Fig. 7, top panel). This contact was lost in the middle of the SMD simulation of recovery from fast inactivation of the apo channel, while the presence of PIP2 and PF-05089771 made it last for almost the whole simulation, showing that the recovery was blocked (Fig. 7, bottom panel). One should note that this simple contact measure does not include the differences in SMD forces. Interestingly, substitution of the residue corresponding to F1552 (F1291 in rNav1.4) was very recently shown to remove a significant amount of fast inactivation, as F1552 possibly forms a T-shaped π–π interaction with F1565 from MFM [18]. This interaction is most probably responsible for coupling the MFM particle to the pore [18].

**Figure 7.**
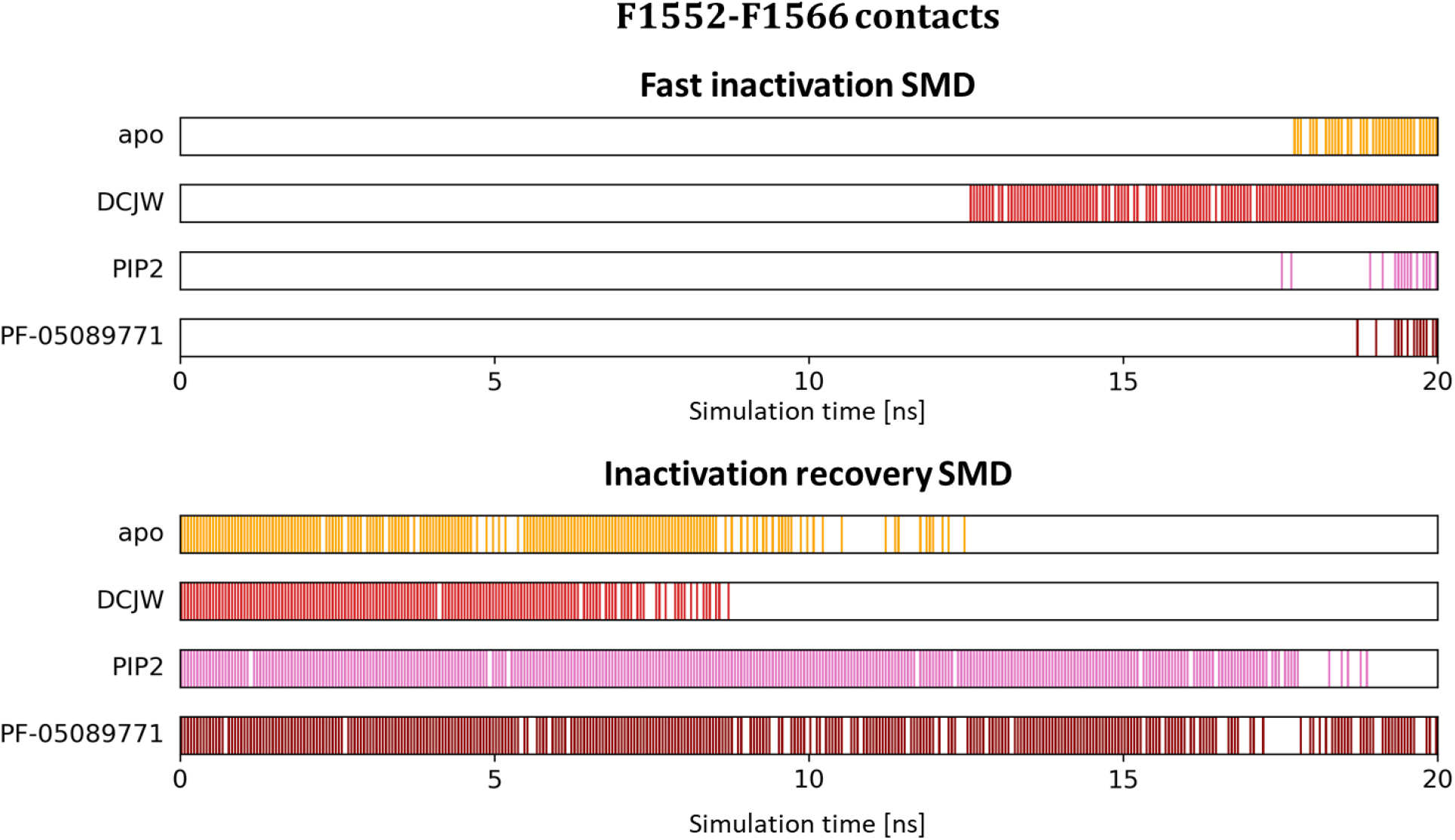
The frequency of the F1565-F1552 contact during SMD simulations of MFM motif binding (fast inactivation, top panel) and unbinding (the recovery from fast inactivation, bottom panel) based on three trajectories of each system.

### 3.6 Experimental part on *P. americana* neuronal preparation

To validate the SMD results, we analyzed the ligands’ impact on the extracellular activity of the semi-isolated cercal nerve of *Periplaneta americana*. The cercal nerve comprises approximately 200 primary sensory neurons and is readily accessible for pharmacological agents.

Since all the tested ligands were dissolved in DMSO, we performed control recordings using 0.1% DMSO in physiological saline. For each recording, we established the basal activity, measured as the average frequency of extracellular action potentials during the 30 seconds preceding the application of the tested solution. We then calculated the percentage change in activity based on the 30-second average frequency of extracellular action potentials recorded 5 minutes after applying the tested solution. 0.1 % DMSO did not influence the frequency of generation of action potentials, and 5 minutes after its application, the activity remained at 98.00 ± 1.59 % (Fig. 8). This confirms that DMSO in 0.1% concentration does not disturb cell membrane integrity, as indicated before [53]. Results for 0.1% DMSO served as a control for further comparisons.

**Figure 8.**
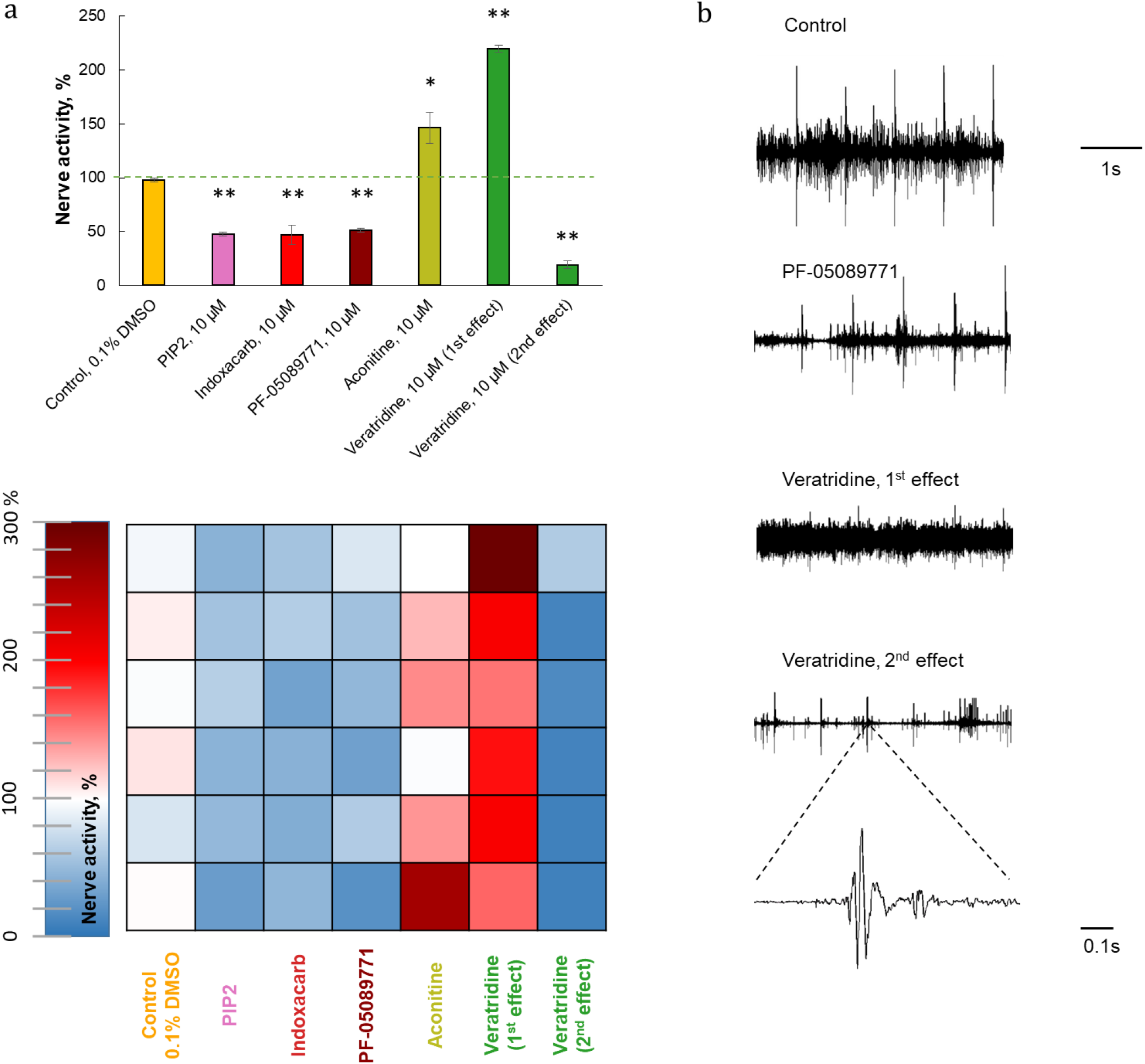
Extracellular recordings of *Periplaneta americana* cercal nerve activity in the presence of various ligands. a) The average nerve activity is expressed as a percentage of basal activity measured before administering the tested solutions. 0.1 % DMSO was used as a solvent for all solutions and a comparison control. Each nerve activity was measured after 5 minutes of incubation with the solutions, except for veratridine, whose effect was observed immediately after drug administration (1^st^ effect) and after 5 minutes of incubation (2^nd^ effect). In the top panel, the data is expressed as mean values ± SE, n = 6. The statistically significant differences between the tested ligand and control (0.1 % DMSO) are marked: ^*^ p < .05, ^**^p < .01. The representation of nerve activity of each recording (n = 6) expressed as a color gradient where blue indicates a decrease in nerve activity and red shows an increase in nerve activity is shown in a bottom panel. c) The representative examples of original recordings of cercal nerve activity for: 1) basal nerve activity; 2) nerve activity 5 minutes after PF-05089771 administration; 3) nerve activity immediately after veratridine 10 µM administration; 4) nerve activity 5 minutes after veratridine 10 µM administration. The bottom panel shows an example of a single extracellular action potential at 0.1 s time base.

All the tested ligands were applied at a 10 µM concentration, which was sufficient to observe their effect and allowed us to compare their potency. First, we tested the Na_v_ activators. Aconitine at 10 µM increased the frequency of action potentials to 146.52 ± 9.23 % (p = 0.045). This corresponds to its well-described effect: aconitine significantly delays the inactivation of Na_v,_ making the neurons more excitable [54]. The action of a more potent activator, 10 µM veratridine, followed a two-step process. Firstly, immediately after its application, veratridine increased cercal nerve activity to 219.92 ± 14.3% (p = 0.005), which also resulted in the loss of response to external stimuli (Fig. 8b). However, nerve activity subsequently decreased in the presence of veratridine, reaching 19.19 ± 3.23% (p = 0.005) five minutes after its application. The two-step effect of veratridine has been described previously [55, 56]; the first effect corresponds to the inhibition of Na_v_ inactivation, which subsequently leads to neuronal membrane depolarization and, as a result, a decrease in nerve activity (the second effect).

Next, we observed how Na_v_ inhibitors change nerve activity. Indoxacarb, a precursor of DCJW, at a concentration of 10 µM, decreased nerve activity to 44.68 ± 1.72% (p = 0.005). DCJW has been reported to keep Na_v_ in its inactivated state [57] and cause neuronal membrane hyperpolarization [58], leading to decreased neuronal activity. Similar observations has been made for 10 μM PF-05089771, which caused a decrease in frequency of action potentials to 51.00 ± 3.41 % (p = 0.008) with a visible reduction of action potentials amplitude (Fig. 8b). PF-05089771 has been shown to enhance inactivation of hNa_v_1.7 channels with the reduction of peak action potential amplitude [59, 60]. Promoting the inactivation state by this drug results in a decrease in cercal nerve activity in our study. Importantly, this is the first report that PF-05089771, primarily developed as an analgesic drug, is also effective on Periplaneta americana Na_v_. The third inhibitor, PIP2, reduced the nerve activity to 47.80 ± 1.87 % (p = 0.005). By accelerating the transition to the inactivated state and slowing recovery from the inactivation of Na_v_ [42], PIP2 decreased neuronal activity in our preparation.

The data obtained from electrophysiological experiments are consistent with the SMD results and, therefore, constitute an experimental confirmation of the results obtained *in silico*.

### 3.7 Limitations

First, investigating the ligand efficacy to affect channel gating requires finding a proper binding position in docking if the experimental data (e.g., cryo-EM structure of a ligand-bound channel) is unavailable. Next, it is not recommended to analyze the whole pathway of conformational changes comprising the fast inactivation process based on SMD simulations analysis, as the addition of forces may lead to non-physiological deformations. While the recorded forces reflect instantaneous resistance to pulling and enable qualitative comparisons between conditions, they should also not be interpreted as free energy barriers (which would require more rigorous methods such as Umbrella Sampling or Metadynamics). Instead, peaks in the force profiles identify critical points where structural rearrangements occur, revealing ligand-dependent stabilization or destabilization of specific functional states. Finally, simulations were performed based on models inferred from homology with the C-terminus, N-terminus, and two long intracellular loops removed. We cannot exclude the possibility that the removed parts could play a role in the fast inactivation process. However, our models were enough to show differences in ligands efficacy in gating modulation and thus to present a new application of the SMD method. Future studies will benefit from the emergence of high-resolution structures of Na_v_ channels with the lacking fragments resolved.

## 4. Conclusions

Here, we propose a simple approach for estimating the potency of ligands or mutations to affect the gating mechanism of Na_v_ channels. The fast inactivation of Na_v_s is a critical step in the channel functional cycle that rapidly terminates each action potential to generate spikes of neuron firings and thus for preventing hyperexcitability. Although binding of the IFM/MFM particle does not describe the whole inactivation process, this step is indispensable in a sequence of conformational changes coupling the voltage-driven movements of VSDIV with the inner mouth of the pore [18]. Moreover, its relocation is easy to track, and – as we show here – the efficacy of known and new chemicals to impair fast inactivation can be assessed by their ability to modify IFM/MFM movement. It allowed us to describe the mode of action of PF-05089771 and PIP2 on insect Na_v_s, which we further validated experimentally by measuring the cockroach neural activity in their presence. Our relatively simple approach appeared to serve well in investigating the various ligands’ impact on Na_v_ gating. After proper system preparation (especially establishing directions of pulling force), it can be used at a mass scale, e.g., as a stage of virtual screening on human channels, followed by electrophysiological recording of ion conductance. Depending on the project objectives, the most promising compounds could be further tested in cell lines with functionally expressed target proteins using the patch clamp technique.

## Data availability

The modified versions of multiSMD, both for IFM/MFM binding and unbinding, together with Python analysis scripts, are available as github repository: https://github.com/kszewc/multiSMD_inactivation. PaNav1 models used here with docked ligands and SMD outputs obtained, together with the scripts to plot the results, are available at RepOD at https://doi.org/10.18150/GLHCMS.

## Acknowledgements

This research was funded by the National Science Centre, Poland, under grant no. 2021/41/N/NZ3/02165 (B.N.). Computations were carried out using the computers of Centre of Informatics Tricity Academic Supercomputer & Network (CI TASK). We also gratefully acknowledge Polish high-performance computing infrastructure PLGrid (HPC Centers: ACK Cyfronet AGH, CI TASK) for providing computer facilities and support within computational grant no. PLG/2024/016996.

## CRediT authorship contribution statement

### Beata Niklas

Conceptualization, Data curation, Formal analysis, Funding acquisition, Investigation, Methodology, Project administration, Visualization, Writing – original draft.

### Milena Jankowska

Investigation, Methodology, Visualization, Writing – review and editing.

### Katarzyna Walczewska-Szewc

Methodology, Software, Validation, Writing – review and editing.

### Bruno Lapied

Supervision, Writing – review and editing.

### Wiesław Nowak

Conceptualization, Supervision, Writing – review and editing.

