## Supplementary Information for "Evaluating the ligands’ potency to modulate the fast inactivation of voltage-gated sodium channel"

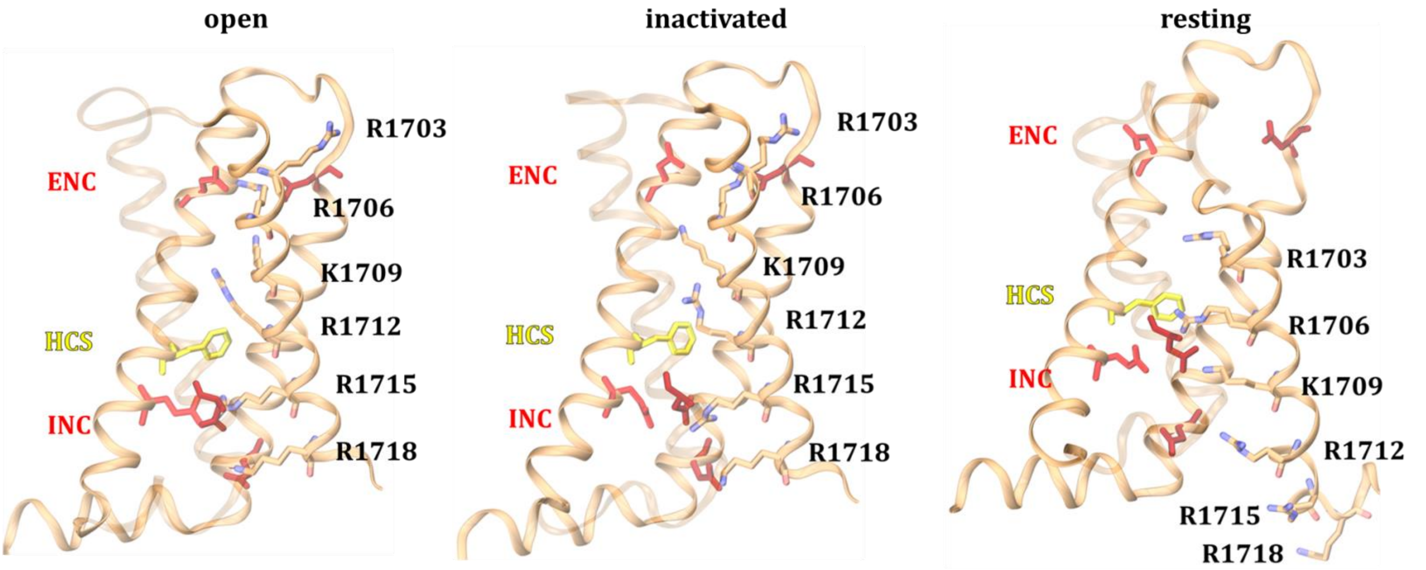


SI Figure 1. The voltage-sensing domain IV (VSDIV) of the *Periplaneta americana* PaNav1 channel in its three main functional states: open (left panel), inactivated (middle panel), and resting (right panel). The gating charges of VSDIV S4 helix are shown as gold sticks with the nitrogen atoms in blue. The residues contributing to the extracellular negative clusters (ENC, red), hydrophobic constriction site (HCS, yellow), and intracellular negative cluster (INC, red) are shown to compare the position of gating charges in three conformational states.

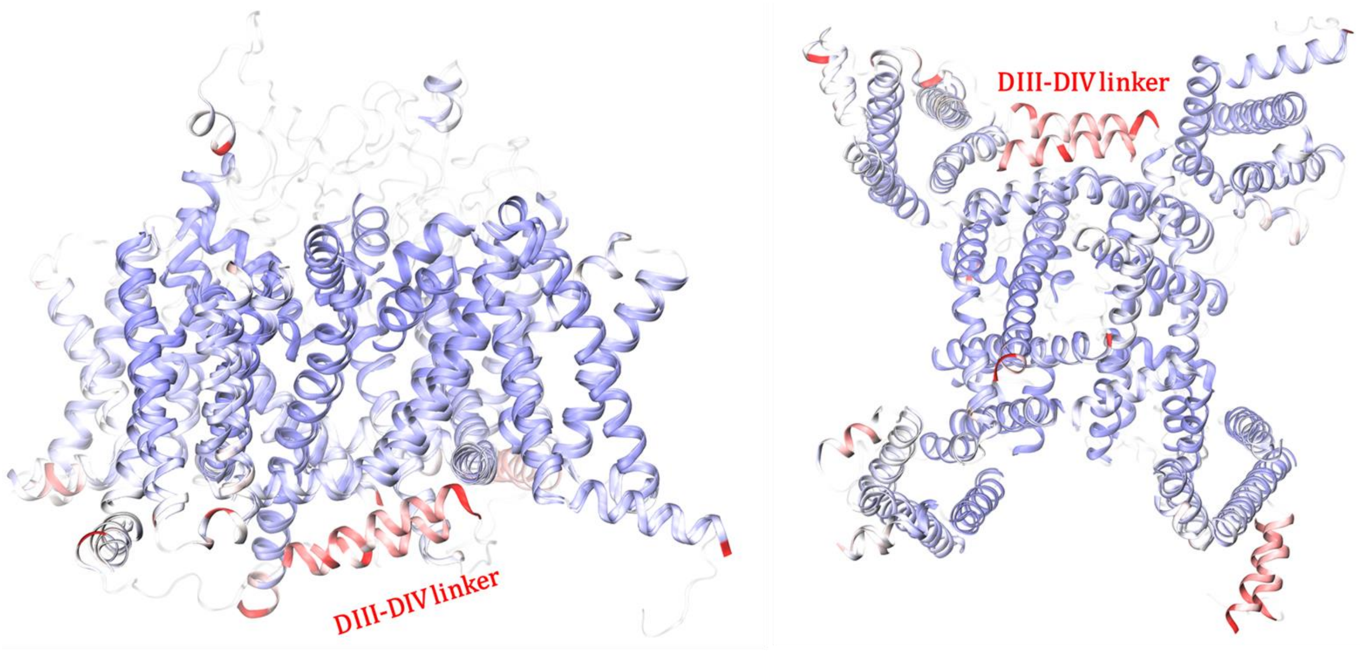


SI Figure 2. The alignment of the open and inactivated-state models showing the main structural differences between them. The regions with the highest structural similarity measured using the root mean square deviation (RMSD) parameter are shown in blue, while the regions exhibiting the greatest structural differences between aligned models are in red. Both extracellular and intracellular loops were excluded from the analysis. The DIII-DIV linker carrying the inactivation particle MFM motif is labeled.

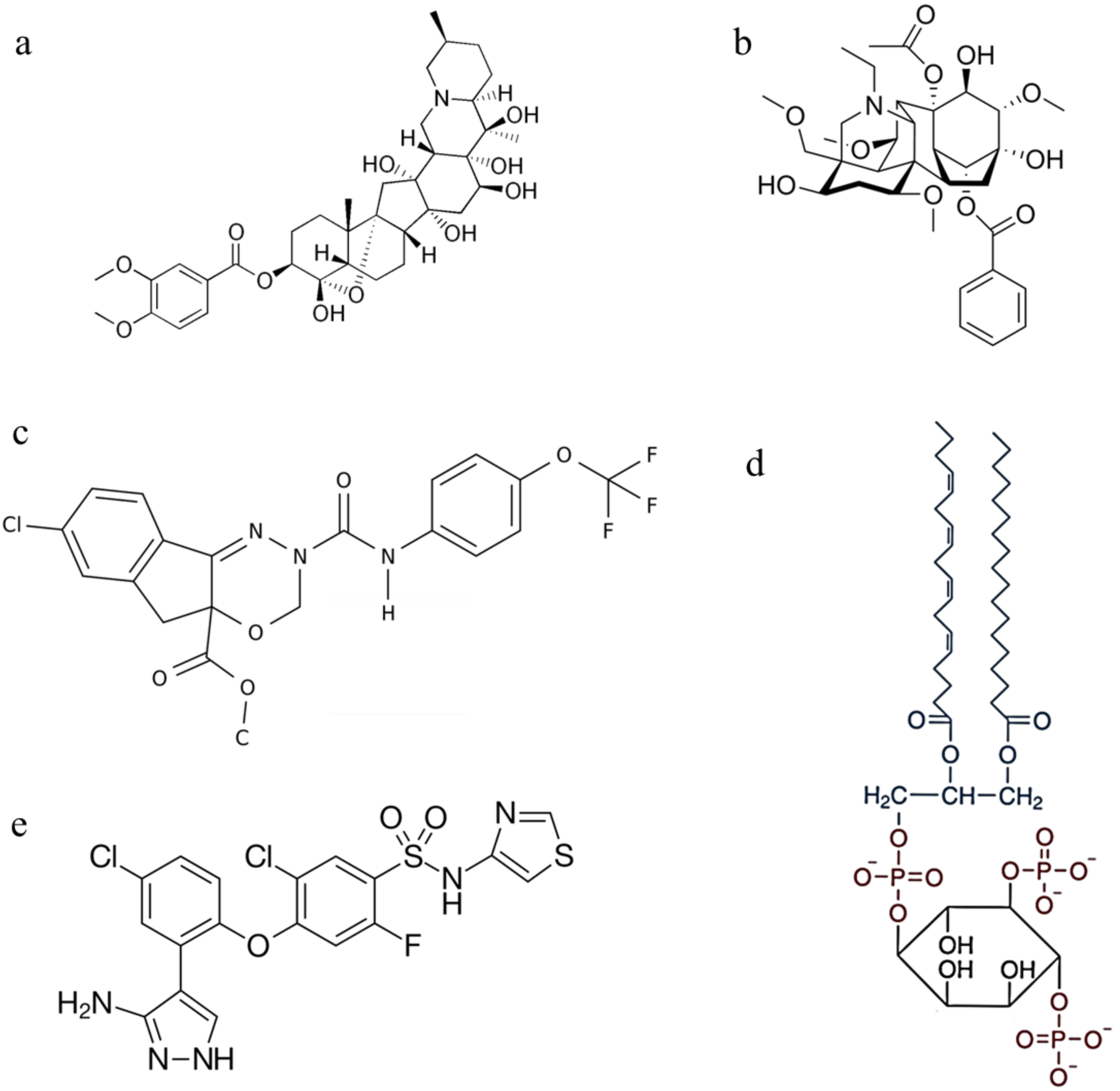


SI Figure 3. Chemical structures of investigated ligands: a) veratridine, b) aconitine, c) N-decarbomethoxyllated JW062 (DCJW), d) phosphatidylinositol 4,5-bisphosphate (PIP2), e) PF-05089771

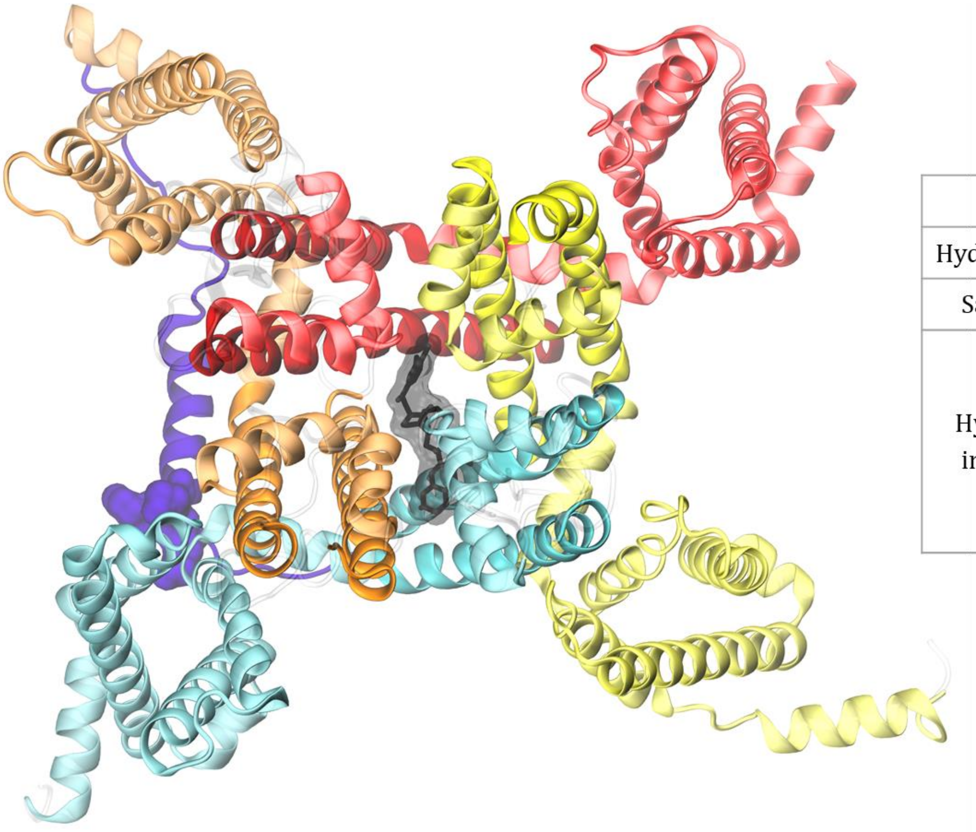

| veratridine interactions | |
|---|---|
| Hydrogen bonds | Ala1495 |
| Salt Bridges | Lys1498 |
| Hydrophobic interactions | Trp1430<br>Phe1492<br>Leu1541<br>Phe1544<br>Val1834<br>Phe1837 |

SI Figure 4. The lowest energy docking pose of veratridine. The inactivated-state model of the PaNav1 channel is shown in the top view (left panel) with four domains colored: DI – red, DII – yellow, DIII – blue, DIV – orange, and the DIII-DIV linker colored violet with the MFM motif marked in a surface representation. The ligand is shown in black in the left panel. The receptor-ligand interactions, found using PLIP server [1], are listed in the right panel.

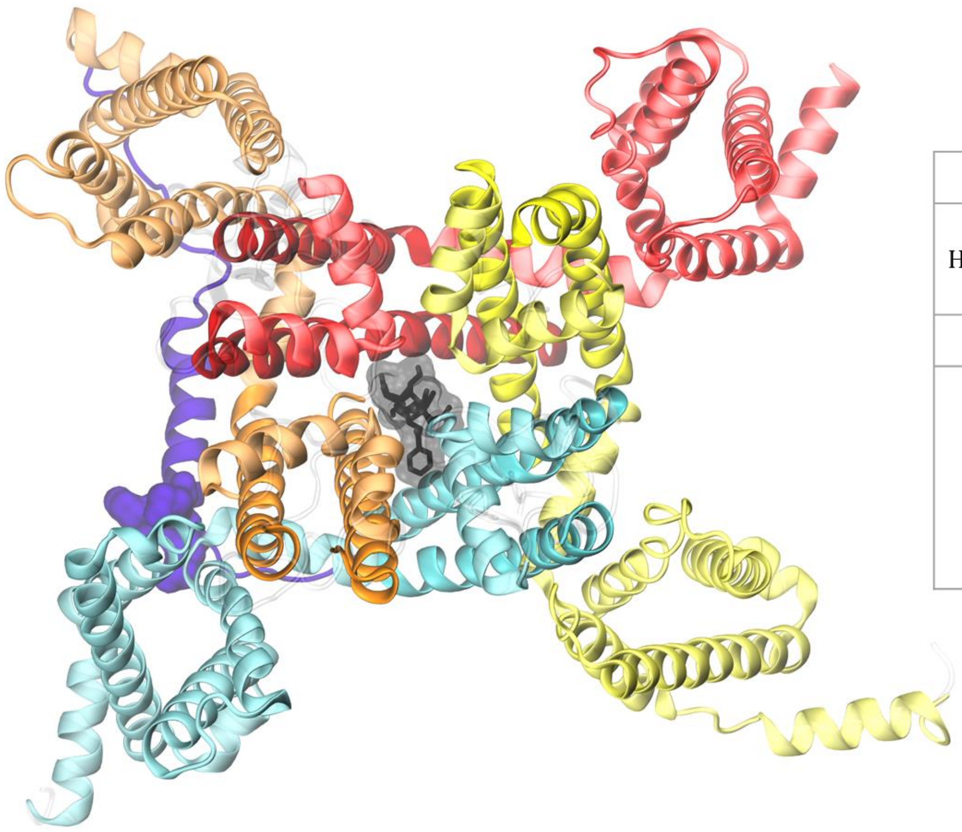

| aconitine interactions | |
|---|---|
| Hydrogen bonds | Phe417<br>Lys1498<br>Thr1789 |
| Salt Bridges | Lys1498 |
| Hydrophobic interactions | Phe417<br>Phe1492<br>Thr1496<br>Leu1541<br>Leu1833<br>Phe1837 |

SI Figure 5. The lowest energy docking pose of aconitine. The inactivated-state model of the PaNav1 channel is shown in the top view (left panel) with four domains colored: DI – red, DII – yellow, DIII – blue, DIV – orange, and the DIII-DIV linker colored violet with the MFM motif marked in a surface representation. The ligand is shown in black in the left panel. The receptor-ligand interactions, found using PLIP server [1], are listed in the right panel.

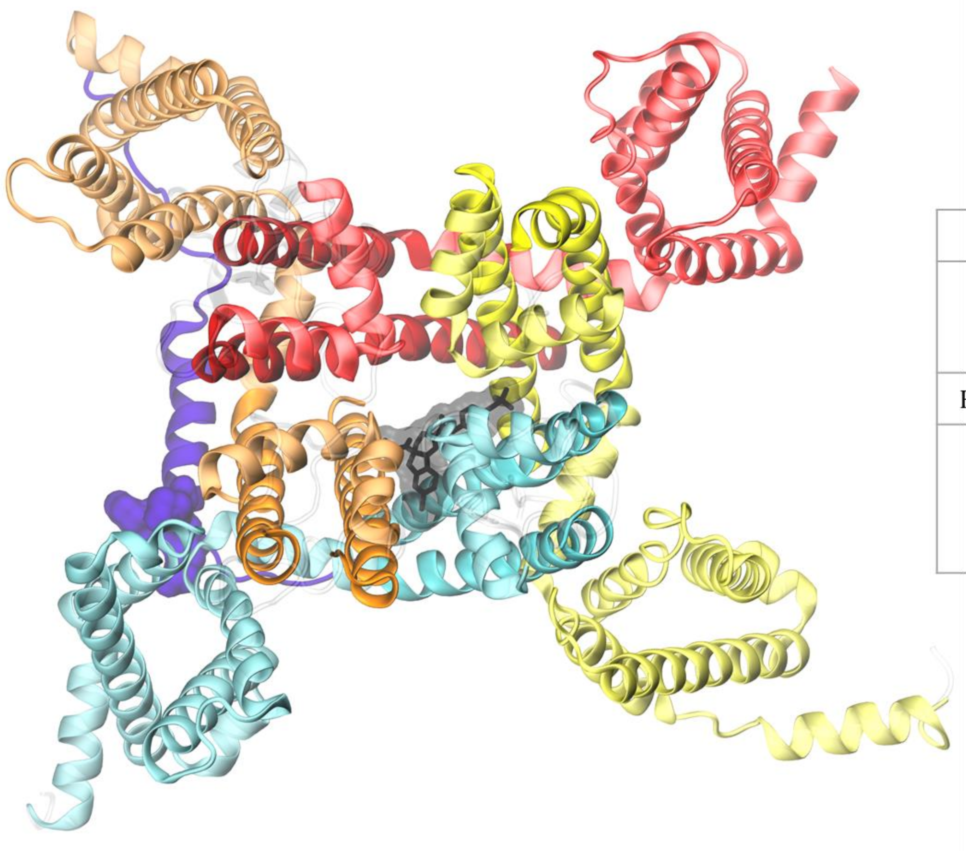

| DCJW interactions | |
|---|---|
| pi-Stacking | Phe417<br>Lys1498<br>Thr1789 |
| Halogen Bonds | Ile1533 |
| Hydrophobic interactions | Phe1492<br>Thr1496<br>Ley1833<br>Phe1837 |

SI Figure 6. The lowest energy docking pose of DCJW. The inactivated-state model of the PaNav1 channel is shown in the top view (left panel) with four domains colored: DI – red, DII – yellow, DIII – blue, DIV – orange, and the DIII-DIV linker colored violet with the MFM motif marked in a surface representation. The ligand is shown in black in the left panel. The receptor-ligand interactions, found using PLIP server [1], are listed in the right panel.

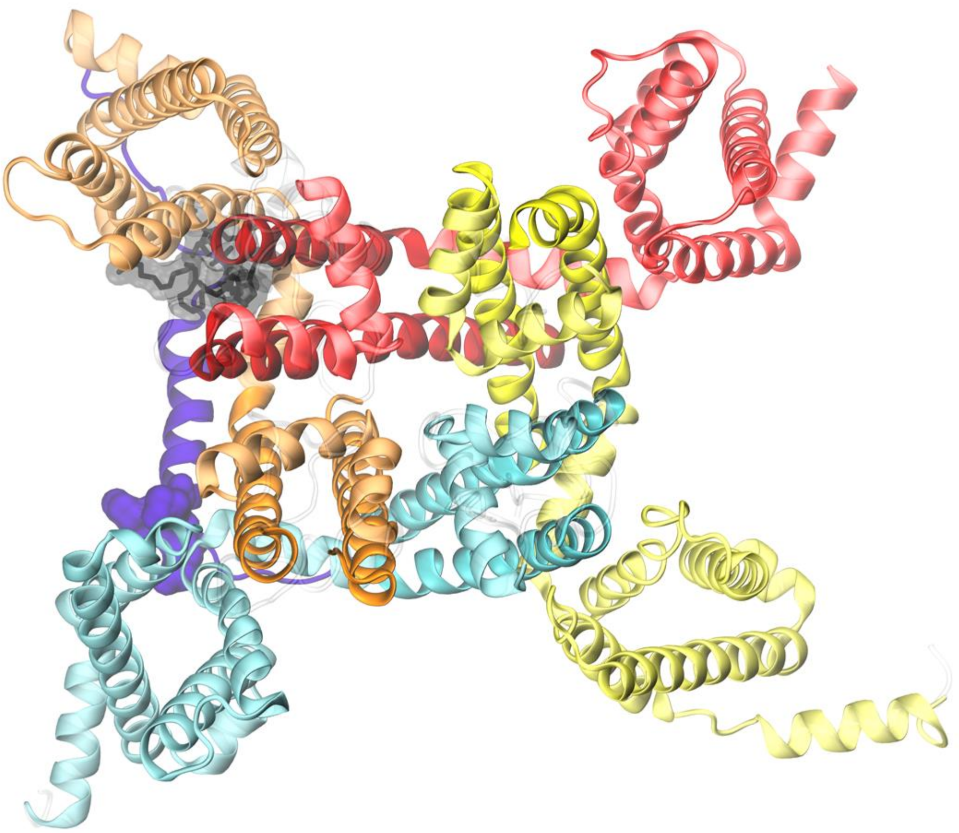

| PIP2 interactions | |
|---|---|
| Hydrogen bonds | Trp1671<br>Arg1724 |
| Salt Bridges | Lys1721<br>Arg1724 |
| Hydrophobic interactions | Phe279<br>Val283<br>Leu414<br>Pro1670<br>Trp1671<br>Leu1673<br>Phe1674<br>Val1677<br>Leu1714<br>Val1717<br>Ile1723<br>Arg1724<br>Leu1727<br>Phe1728 |

SI Figure 7. The lowest energy docking pose of phosphatidylinositol 4,5-bisphosphate (PIP2). The inactivated-state model of the PaNav1 channel is shown in the top view (left panel) with four domains colored: DI – red, DII – yellow, DIII – blue, DIV – orange, and the DIII-DIV linker colored violet with the MFM motif marked in a surface representation. The ligand is shown in black in the left panel. The receptor-ligand interactions, found using PLIP server [1], are listed in the right panel.

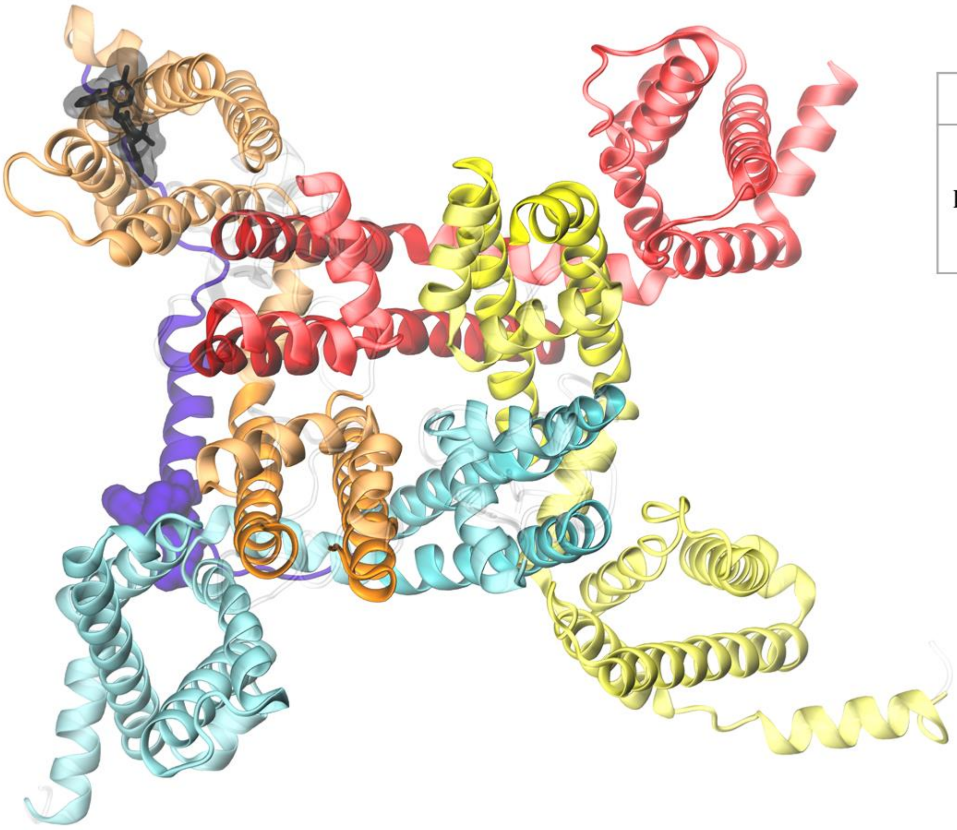

| PF-05089771 interactions | |
|---|---|
| Hydrogen bonds | Asn1644<br>Ser1689<br>Asp1690<br>**Arg1706** |

MD-refinement

| PF-05089771 interactions | |
|---|---|
| Hydrogen bonds | Ser1689<br>Asp1690<br>**Arg1712** |
| Halogen Bonds | Asp1638<br>Asp1641 |
| Hydrophobic interactions | Asp1641<br>Tyr1642<br>Ile1648<br>Leu1686 |

SI Figure 8. The lowest energy docking pose of PF-05089771 sulfonamide inhibitor. The inactivated-state model of the PaNav1 channel is shown in the top view (left panel) with four domains colored: DI – red, DII – yellow, DIII – blue, DIV – orange, and the DIII-DIV linker colored violet with the MFM motif marked in a surface representation. The ligand is shown in black in the left panel. The receptor-ligand interactions, found using PLIP server [1], are listed in the right top panel. The additional step of finding receptor-ligand interactions was performed after the equilibration and a 1 ns-long molecular dynamics (MD) simulation (right bottom panel). The hydrogen bond between the ligand and one of the gating charges was found in both cases (bold in tables). The hydrogen bond was previously observed between a sulfonamide inhibitor and the residue corresponding to Arg1706 [2] and Arg1712 [3].

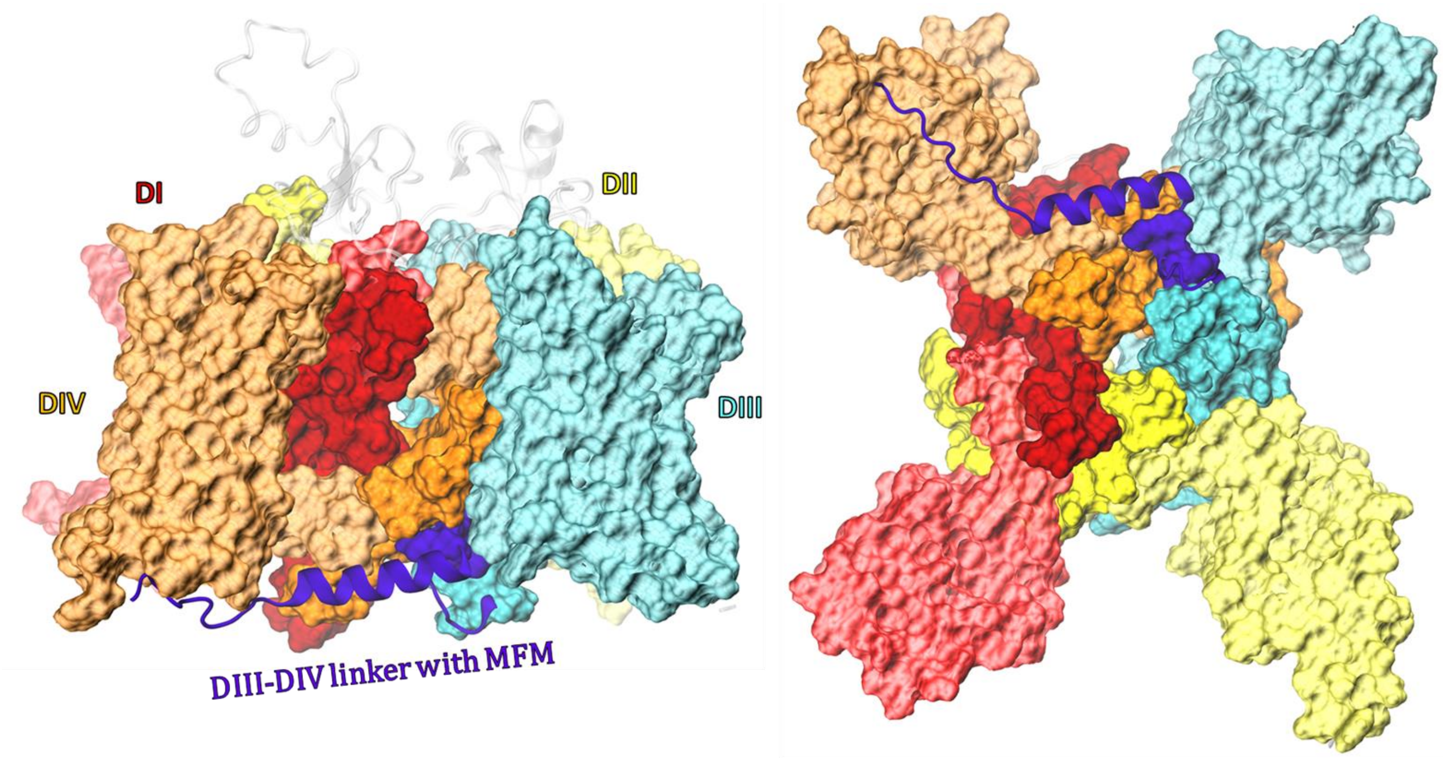


**MFM particle (M1565, F1566, M1567) interactions**

| DIII | DIII-DIV linker | DIV |
|---|---|---|
| V1407 | L1563 | M1732 |
| V1408 | E1564 | S1733 |
| A1411 | T1568 | P1735 |
| L1412 | Q1571 | A1736 |
| F1552 | K1572 | N1739 |
| Q1555 | Y1575 | I1740 |
| K1556 | | N1842 |
| A1559 | | I1845 |
| | | A1846 |
| | | E1850 |

SI Figure 9. The inactivation particle binding pocket. The inactivated-state model of the PaNav1 channel is shown in the side view (top left panel) and the bottom view (top right panel) with four domains in a surface representation, colored: DI – red, DII – yellow, DIII – blue, DIV – orange. The DIII-DIV linker is in violet (here, a surface representation is not applied for clarity) with the inactivation particle MFM motif shown in a surface representation. The list of interactions between residues contributing to the MFM particle (M1565, F1566, and M1567) with its binding pocket is presented in the bottom panel.

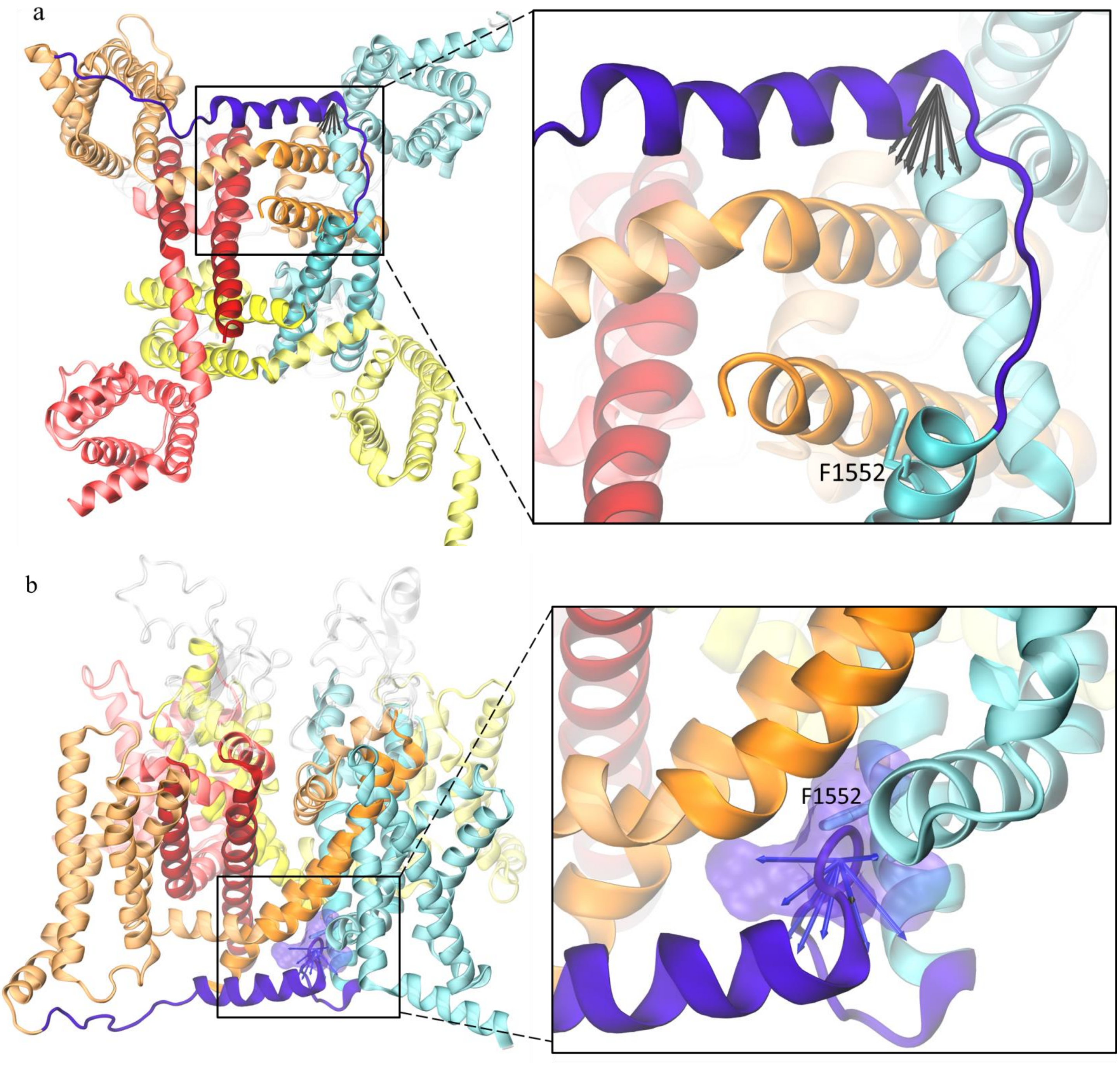


SI Figure 10. The pulling vectors in SMD simulations of the PaNav1 channel. a) The bottom view of the open state model with the pulling vectors shown in black. b) The side view of the inactivated state model with the pulling vectors in blue and the MFM particle occupying its binding pocket shown in the surface representation. F1552 is shown in both panels.

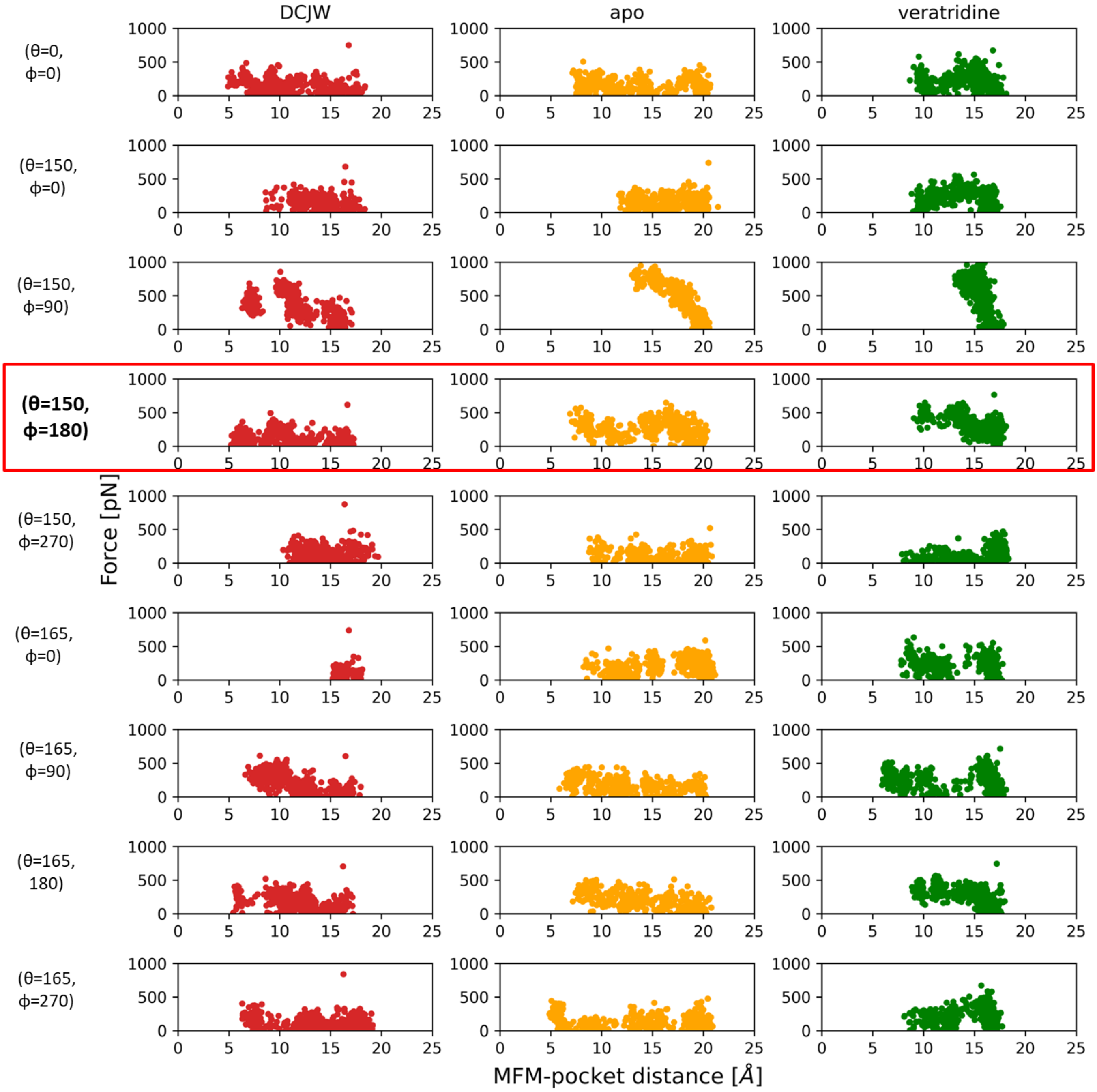


SI Figure 11. The force-distance plots of the MFM particle binding to its pocket based on 20 ns-long SMD simulations of the PaNav1 channel without a ligand (apo, orange), and the ligand-bound channel: DCJW-bound (red) and veratridine-bound (green). Here, nine pulling directions were sampled to choose the one that provides the best representation of MFM unbinding by generating a set of vectors characterized by small variations in theta (0°, 150°, 165°) and phi (0°, 90°, 180°, 270°) angles from the main pulling direction within spherical coordinates. The change in distance between the centers of masses of the MFM particle and its binding pocket (x axis) and the forces taken in simulations (y axis) are presented for all tested pulling directions. The open-state model was used as a starting point.

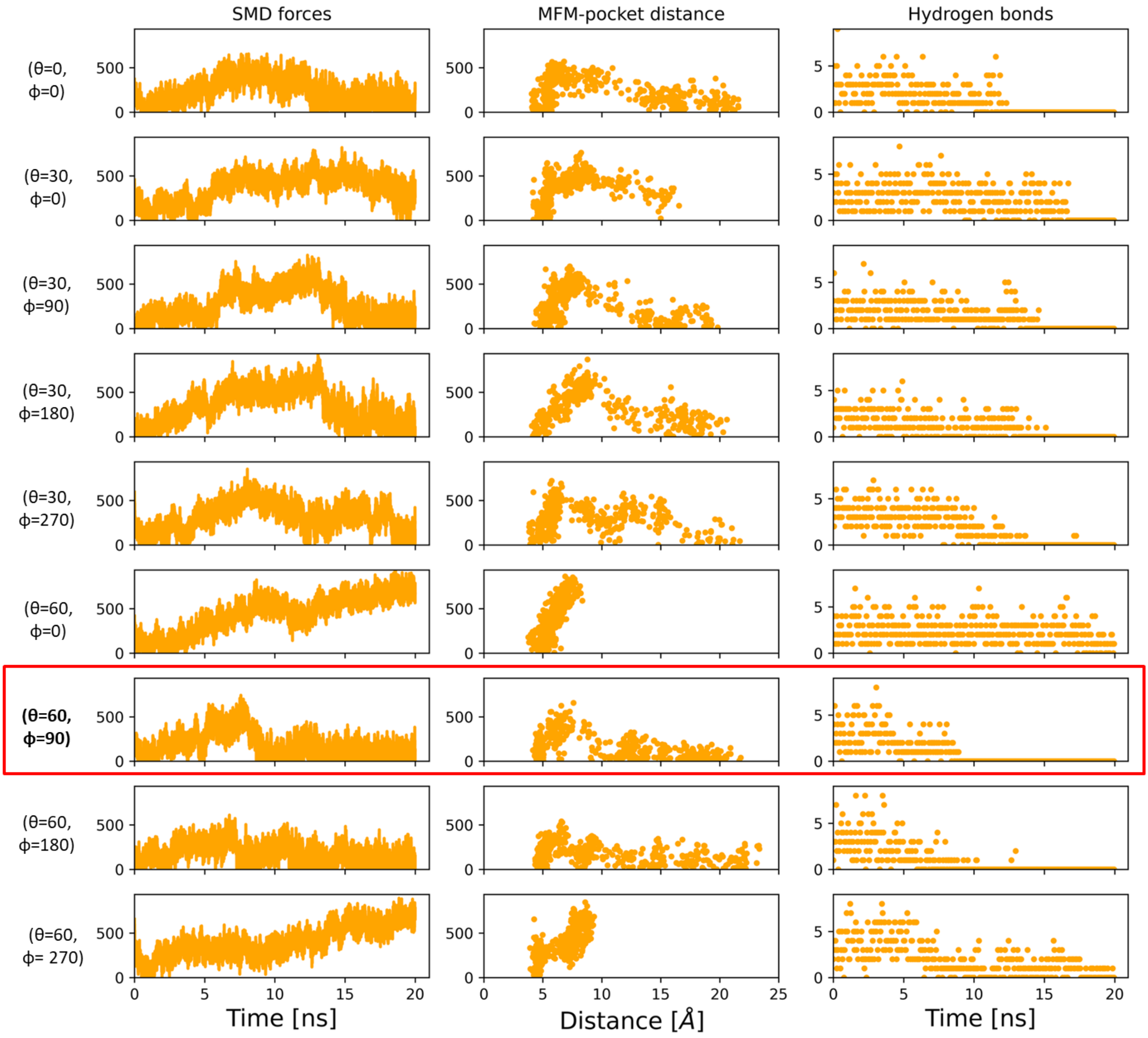


SI Figure 12. The MFM particle displacement from its pocket starting from the inactivated-state PaNav1 model with no ligand, based on SMD simulation (20 ns-long each). Nine pulling directions: theta (0°, 30°, 60°) and phi (0°, 90°, 180°, 270°) were tested. Here, three parameters were investigated to choose the best pulling direction: force vs. time (left panel), distance vs. force (middle panel), and the disruption of hydrogen bonds between F1566 (from MFM) and the binding pocket (right panel). The θ=60°, φ=90° was chosen due to the complete loss of hydrogen bond and the MFM displacement of >20 Å that required the smallest energy.

### Supplementary Information References:

[1] M.F. Adasme, K.L. Linnemann, S.N. Bolz, F. Kaiser, S. Salentin, V.J. Haupt, M. Schroeder, PLIP 2021: Expanding the scope of the protein–ligand interaction profiler to DNA and RNA, Nucleic acids research 49(W1) (2021) W530-W534.
[2] Q. Wu, J. Huang, X. Fan, K. Wang, X. Jin, G. Huang, J. Li, X. Pan, N. Yan, Structural mapping of Nav1. 7 antagonists, Nature communications 14(1) (2023) 3224.
[3] S. Ahuja, S. Mukund, L. Deng, K. Khakh, E. Chang, H. Ho, S. Shriver, C. Young, S. Lin, J. Johnson Jr, Structural basis of Nav1. 7 inhibition by an isoform-selective small-molecule antagonist, Science 350(6267) (2015) aac5464.
